# Homeostatic responses regulate selfish mitochondrial genome dynamics in *C. elegans*

**DOI:** 10.1101/050930

**Authors:** Bryan L. Gitschlag, Cait S. Kirby, David C. Samuels, Rama D. Gangula, Simon A. Mallal, Maulik R. Patel

**Affiliations:** Department of Biological Sciences, Vanderbilt University, Nashville, TN, USA; Interdisciplinary Graduate Program, Vanderbilt University, Nashville, TN, USA; Biological Sciences Graduate Program, Vanderbilt University, Nashville, TN, USA; Department of Molecular Physiology and Biophysics, Vanderbilt University, Nashville, TN, USA; Vanderbilt Genetics Institute, Vanderbilt University, Nashville, TN, USA; Department of Medicine, Vanderbilt University School of Medicine, Nashville, TN, USA; Department of Pathology, Microbiology and Immunology, Vanderbilt University School of Medicine, Nashville, TN, USA; Institute for Immunology and Infectious Diseases, Murdoch University, Murdoch, Western Australia, Australia.; Department of Cell and Developmental Biology, Vanderbilt University, Nashville, TN USA

## Abstract

Selfish genetic elements have profound biological and evolutionary consequences. Mutant mitochondrial genomes (mtDNA) can be viewed as selfish genetic elements that persist in a state of heteroplasmy despite having potentially deleterious consequences to the organism. We sought to investigate mechanisms that allow selfish mtDNA to achieve and sustain high levels. Here, we establish a large 3.1kb deletion bearing mtDNA variant *uaDf5* as a *bona fide* selfish genome in the nematode *Caenorhabditis elegans*. Next, using droplet digital PCR to quantify mtDNA copy number, we show that *uaDf5* mutant mtDNA replicates in addition to, not at the expense of, wildtype mtDNA. These data suggest existence of homeostatic copy number control for wildtype mtDNA that is exploited by *uaDf5* to ‘hitchhike’ to high frequency. We also observe activation of the mitochondrial unfolded protein response (UPR^mt^) in animals with *uaDf5*. Loss of UPR^mt^ results in a decrease in *uaDf5* frequency whereas constitutive activation of UPR^mt^ increases *uaDf5* levels. These data suggest that UPR^mt^ allows *uaDf5* levels to increase. Interestingly, the decreased *uaDf5* levels in absence of UPR^mt^ recover in parkin mutants lacking mitophagy, suggesting that UPR^mt^ protects *uaDf5* from mitophagy. We propose that cells activate two homeostatic responses, mtDNA copy number control and UPR^mt^, in *uaDf5* heteroplasmic animals. Inadvertently, these homeostatic responses allow *uaDf5* levels to be higher than they would be otherwise. In conclusion, our data suggest that homeostatic stress response mechanisms play an important role in regulating selfish mitochondrial genome dynamics.

## Significance statement

Hundreds to thousands of wildtype mitochondrial genomes (mtDNA) are normally present in each cell. However, a fraction of these mtDNA can acquire mutations and coexist with wildtype mtDNA in a state of heteroplasmy. Mutant mtDNA that persist despite having potentially deleterious consequences can be viewed as selfish genetic elements. We sought to determine mechanisms underlying selfish mtDNA dynamics. We show that a stably propagating deletion-bearing mutant mtDNA in *C. elegans* behaves like a selfish genetic element. Presence of mutant mtDNA at high levels activates two stress responses. We further present data suggesting that the mutant mtDNA takes advantage of these responses to persist and proliferate. Our work identifies molecular processes that are important regulators of selfish mitochondrial genome dynamics.

## Introduction

Many ecological systems are predicated upon mutualistic interactions between different organisms (Momeni, Chen et al. 2011). Similar interactions between different genetic entities can underlie fundamental aspects of cell biology. For instance, the nuclear and the mitochondrial genome (mtDNA) work in a coordinated fashion to ensure optimal cellular fitness. However, just like in ecology, mutualistic interactions between these genomes are fraught with the risk that selfish mtDNA could arise. Selfish mtDNA can be defined as mutant mtDNA variants that have a transmission advantage over wildtype mtDNA despite jeopardizing cellular fitness and even organismal survival (Clark, Howe et al. 2012). Such selfish mtDNA variants often replicate and are co-transmitted along with wildtype mtDNA in a state of heteroplasmy (Taylor, Zeyl et al. 2002). The presence of such mutant mtDNA at frequencies above a critical threshold can be pathogenic and is thought to be one of the causes underlying inherited and age-related degeneration and metabolic diseases (Wallace and Chalkia 2013, Stewart and Chinnery 2015).

Selfish mtDNA are most studied in yeast, where they arise at very high rates and result in formation of ‘petite’ colonies with mitochondrial respiration failure (Williamson 2002, Bernardi 2005). Despite harboring major deletions and rearrangements, and causing severe cell growth defects, many of these mutant mtDNA are able to outcompete wildtype mtDNA in yeast cells and are hence dubbed ‘hypersuppressive’ mtDNA (MacAlpine, Kolesar et al. 2001, Harrison, MacLean et al. 2014, Jasmin and Zeyl 2014). While these selfish mtDNA were at first thought to outcompete wildtype mtDNA by virtue of possessing more origins of replication, this interpretation has been questioned in light of new data (Williamson 2002). In metazoans, mutant mtDNA are largely characterized by deletions rather than the massive genomic rearrangements observed in hypersuppressive mtDNA. These data suggest that the mechanisms employed by selfish mtDNA in yeast are fundamentally different than those in metazoans. A deletion-harboring mtDNA was recently discovered in natural populations of the nematode species *C. briggsae*. Interestingly, this mutant mtDNA variant is found across *C. briggsae* strains of diverse geographic origins, suggesting that it has stably persisted on evolutionary time-scales (Howe and Denver 2008, Clark, Howe et al. 2012). The presence of this mtDNA variant decreases fecundity and pharyngeal pumping rates, suggesting that it negatively impacts organismal fitness (Estes, Coleman-Hulbert et al. 2011). While this mutant mtDNA was shown to have a transmission advantage over wildtype mtDNA in mutation accumulation lines and in small populations, the cellular and molecular bases underlying its competitive success are not well understood (Clark, Howe et al. 2012, Phillips, Coleman-Hulbert et al. 2015).

In this study, we establish a mutant mtDNA variant called *uaDf5* as a selfish genetic element in *C. elegans. uaDf5* coexists as a heteroplasmy with wildtype mtDNA despite being slightly deleterious. Next, using droplet digital PCR (ddPCR) to quantify mtDNA copy number dynamics, we show the existence of homeostatic copy number control for wildtype mtDNA. Our data on *uaDf5* mtDNA copy number dynamics are most consistent with *uaDf5* exploiting this mtDNA copy number control to ‘hitchhike’ to high frequency. Finally, we observe activation of the mitochondrial unfolded protein response (UPR^mt^) in *uaDf5* heteroplasmic animals. Loss of UPR^mt^ results in a decrease in *uaDf5* levels, while constitutive activation leads to its increase. Hence, besides mtDNA copy number control, *uaDf5* also exploits UPR^mt^, together suggesting that homeostatic cellular processes are important determinants of selfish mtDNA dynamics.

## Results

### A *bona fide* ‘selfish’ mtDNA in *C. elegans*

We focused on identifying potentially ‘selfish’ mtDNA in a genetically tractable metazoan. *C. elegans* is an ideal model system to study mtDNA dynamics in a multicellular organism. Besides offering a powerful genetic toolkit, *C. elegans* mtDNA shares many conserved features with its mammalian counterpart (Okimoto, Macfarlane et al. 1992). First, like mammalian mtDNA, mtDNA in *C. elegans* is uniparentally inherited through the oocyte (Tsang and Lemire 2002, Al Rawi, Louvet-Vallee et al. 2011, Sato and Sato 2011). Second, *C. elegans* mtDNA encodes 12 of the 13 proteins encoded by mammalian mtDNA. Finally, like in mammals but in contrast to yeast, most large-scale mutations in *C. elegans* mtDNA are deletions rather than genomic rearrangements (Denver, Morris et al. 2000).

Previous studies have identified a *C. elegans* strain harboring mutant mtDNA called *uaDf5*, which has a 3.1kb deletion that removes four protein-coding genes and seven tRNAs (Fig. 1A) (Tsang and Lemire 2002). Due to this large deletion, individuals carrying only *uaDf5* mtDNA would not be expected to be viable. Indeed, animals homoplasmic for *uaDf5* have not been reported (Tsang and Lemire 2002, Liau, Gonzalez-Serricchio et al. 2007). Remarkably however, animals that have lost *uaDf5* have also not been previously observed, even after passaging over hundreds of generations (Tsang and Lemire 2002, Liau, Gonzalez-Serricchio et al. 2007). Moreover, *uaDf5* levels steadily increase in individuals that inherit it at a low frequency (Tsang and Lemire 2002).

**Figure 1.**
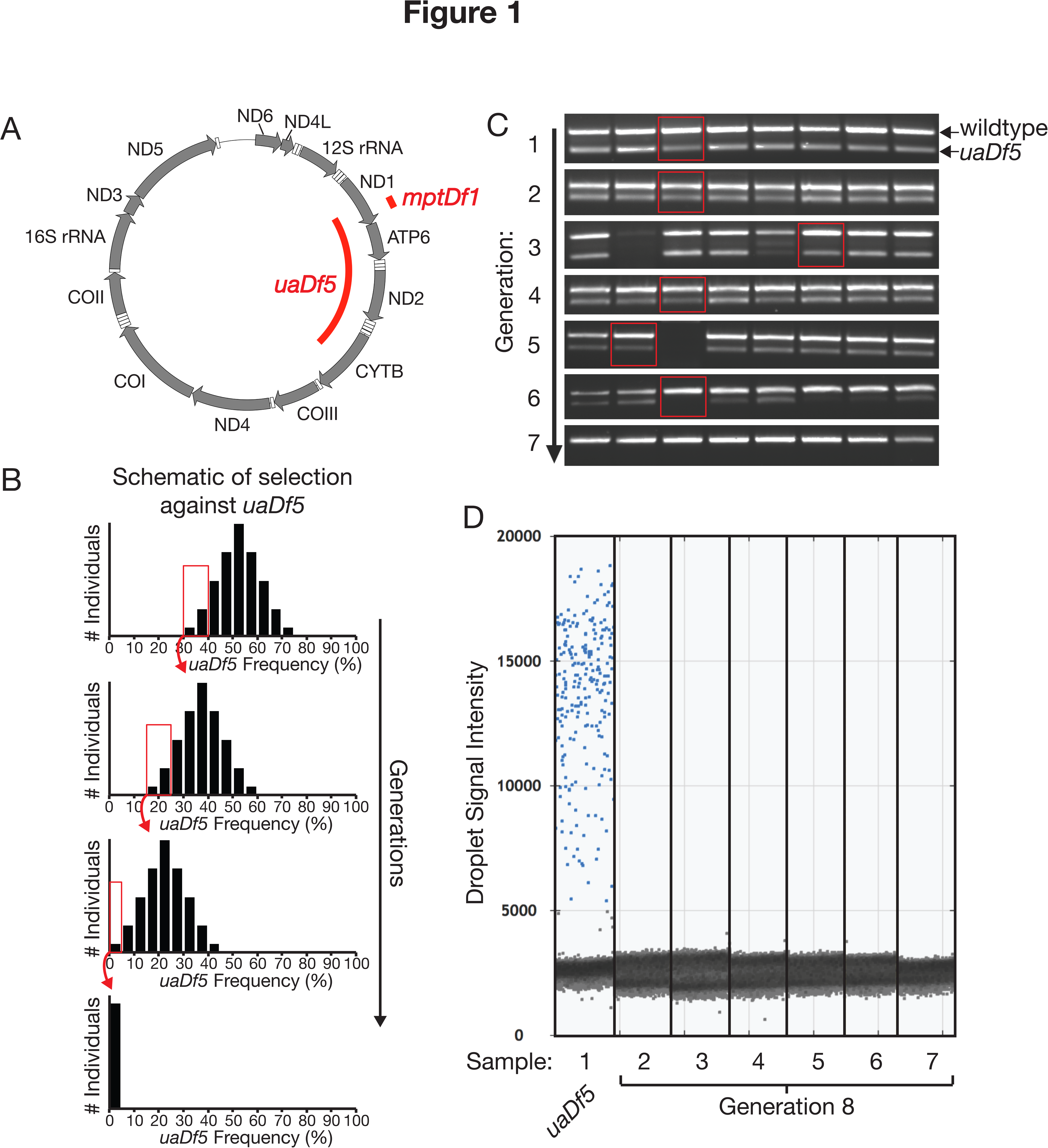
Mutant mtDNA*uaDf5* can be forced out from a stably persisting heteroplasmy in *C. elegans*. (A) Schematic of *C. elegans* mtDNA showing the *uaDf5* and *mptDf1* deletions (long and short red bars, respectively). Grey arrows show protein and rRNA-encoding genes and their orientation. White boxes show genes encoding tRNAs. (B) Schematic illustrating the selection strategy to force loss of *uaDf5* mtDNA from a heteroplasmic *C. elegans* line. Each generation, the progeny of individuals with the lowest *uaDf5* levels were selected for subsequent propagation. (C) Single worm PCR of wildtype and *uaDf5* mtDNA. Successive propagation of individual worms with low *uaDf5* levels (red boxes) results in complete loss of *uaDf5* mtDNA from the population over multiple generations. (D) ddPCR data from single worms confirming complete loss of *uaDf5*. Positive droplets containing *uaDf5*-specific PCR product exhibit increased fluorescence intensity (blue) compared to negative droplets that contain no *uaDf5* mtDNA (gray). For each droplet, the droplet reader detects droplet size, shape, and fluorescence intensity, and automatically distinguishes positive from negative droplets on the basis of these criteria. Sample 1, control containing *uaDf5*.

One explanation for the stable maintenance of *uaDf5* over many generations is balanced heteroplasmy, in which two mtDNA haplotypes possess lethal but non-overlapping mutations. In this scenario, neither mtDNA type can be lost because neither mtDNA is capable of fully supporting viability. In order to test this hypothesis, we sought to sequence the entire non-*uaDf5* mtDNA in the heteroplasmic animals. By designing primers inside the *uaDf5* deletion, we were able to specifically amplify the entire genic region of non-*uaDf5* mtDNA as two large PCR products (Fig. S1). Under the ‘balanced heteroplasmy’ hypothesis, we expected to find deleterious mutations in the non-*uaDf5* mtDNA in the *uaDf5* heteroplasmic individuals. In contrast, upon sequencing, we found that the non-*uaDf5* mtDNA in *uaDf5* heteroplasmic animals is wildtype. Next, to sequence an approximately 500bp highly AT-rich non-coding region that was not captured within the two PCR products, we amplified just the non-coding region of the mtDNA using primers that are common to both *uaDf5* and non-*uaDf5* mtDNA. Upon sequencing however, we did not observe any apparent heteroplasmic mutations, as we might expect if there were any mutations specific to the non-*uaDf5* mtDNA. Thus, our sequencing data do not support balanced heteroplasmy as an explanation for *uaDf5* persistence since the non-*uaDf5* mtDNA is wildtype, suggesting that *uaDf5* is not critical for organismal viability.

**Figure S1.**
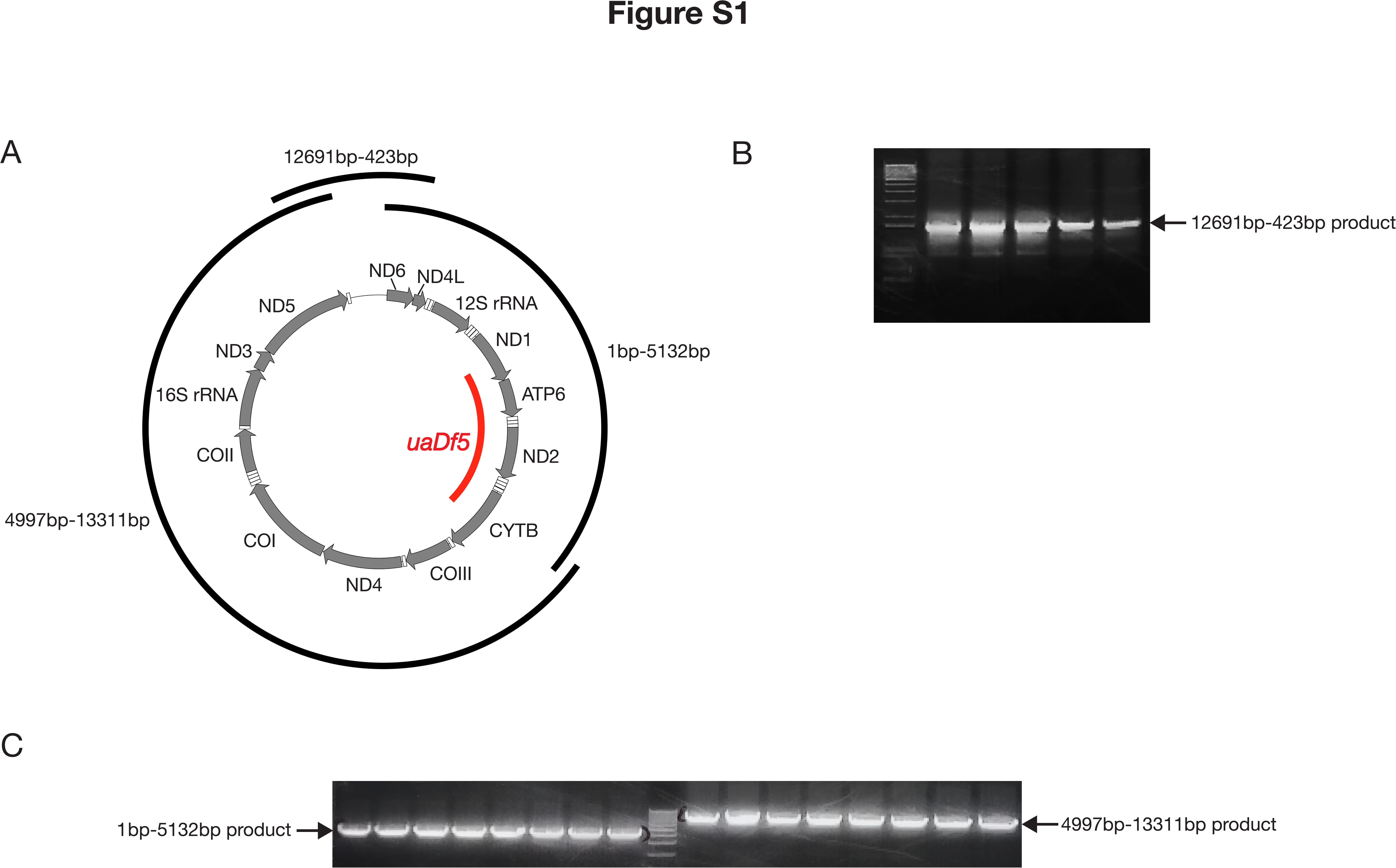
Three PCR products used for sequencing of non*uaDf5* mtDNA, mapped to *C. elegans* reference mtDNA. (A) Three overlapping PCR products were amplified from *C. elegans* mtDNA. The reverse primer used for generating the PCR product spanning base pair positions 1 through 5,132 is specific to a region within the *uaDf5* deletion, as is the forward primer used for generating the PCR product spanning base pair positions 4,997 through 13,311. Additional primers were used for sequencing all three PCR products (see Materials and Methods). (B) Gel image of product amplified by PCR across the D-Loop region. (C) Gel image of products amplified by PCR across the protein-coding mtDNA regions.

Our sequencing data are instead consistent with *uaDf5* behaving like a selfish genetic element. We reasoned that if *uaDf5* is not critical for viability, then we should be able to recover healthy individuals that have lost it. In order to test this hypothesis, we carried out additional experiments in which we selected individuals with progressively lower *uaDf5* levels across multiple generations (red boxes in schematic in Fig. 1B). Under this ‘selection’ regime, we were able to recover healthy individuals homoplasmic for wildtype mtDNA that had undetectable levels of *uaDf5* (Fig. 1C), suggesting that these individuals have lost *uaDf5* mutant mtDNA. To confirm complete loss of *uaDf5*, we conducted droplet digital PCR (ddPCR) with *uaDf5* specific primers. ddPCR relies on performing thousands of independent PCR reactions simultaneously in small droplets that are designed to hold an average of one template per droplet (Hindson, Ness et al. 2011). Hence, it provides a highly sensitized way to detect rare variants. We observed complete loss of *uaDf5* using ddPCR (Fig. 1D). Together with our sequencing results, our finding that *uaDf5* can be eliminated rules out balanced heteroplasmy as an explanation for *uaDf5* maintenance. Instead, together with the previously published data that *uaDf5* has a transmission advantage when inherited at low frequencies (Tsang and Lemire 2002) and is associated with decreased organismal fitness (Liau, Gonzalez-Serricchio et al. 2007), we consider the second alternative that *uaDf5* is a selfish mtDNA.

### *uaDf5* has ‘runaway’ copy number dynamics while wildtype mtDNA levels are tightly controlled

Although *uaDf5* mtDNA levels are variable across individuals, previous reports have suggested they can reach high frequency in individuals (Tsang and Lemire 2002). Indeed, we were able to stably maintain laboratory populations in which individual worms are heteroplasmic for *uaDf5* mtDNA, where *uaDf5* comprises 60-80% of overall mtDNA present in each worm (Fig. 2A). To ascertain mechanisms that regulate *uaDf5* levels, we next examined how levels of mutant *uaDf5* mtDNA affected the levels of wildtype mtDNA. One simple ‘direct competition’ model suggests that wildtype mtDNA levels should be inversely proportional to *uaDf5* mtDNA levels, while total mtDNA levels remain consistent. Alternatively, we might find that the copy number of wildtype mtDNA is maintained independent of *uaDf5* levels.

**Figure 2.**
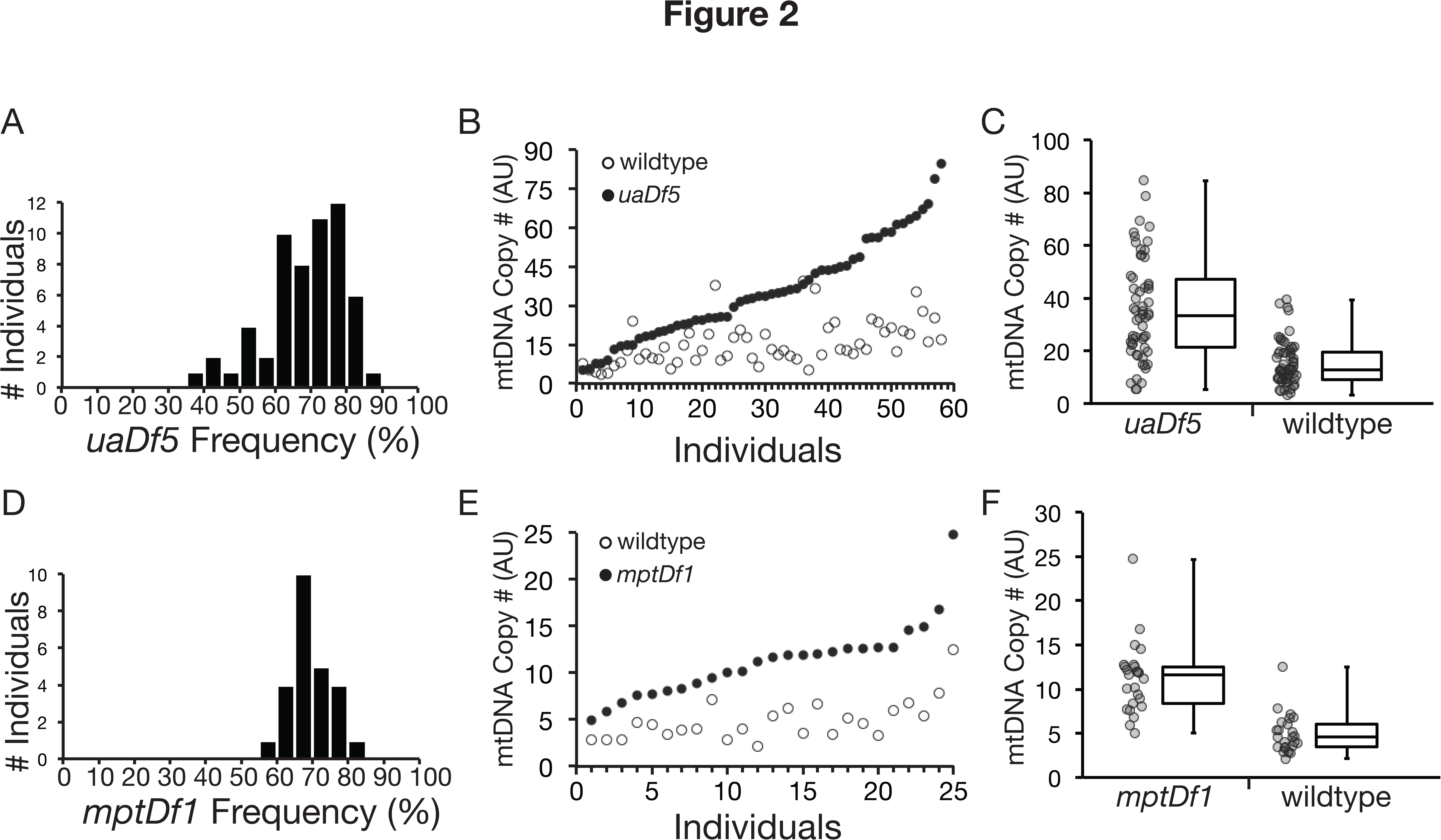
Quantification of mtDNA copy number dynamics reveals mtDNA copy number control. (A) Histogram showing *uaDf5* frequency (%) distribution in individuals from a population stably maintaining *uaDf5* mtDNA (n=58). Heteroplasmy frequency was determined using ddPCR to quantify wildtype and *uaDf5* mtDNA copy number in single individuals. (B) mtDNA levels in individual day 4 adult worms, normalized to actin and rank-ordered by *uaDf5* mtDNA copy number. (C) Wider variation in *uaDf5* relative to wildtype copy number (p<0.05) suggests that wildtype mtDNA, but not *uaDf5* mtDNA, is subject to homeostatic copy number control. Grey data points show mtDNA copy number from single individuals. Box and whisker plot shows the median, lower and upper quartile (boxes), and minimum and maximum (error bars) mtDNA copy number. (D) *mptDf1* frequency distribution obtained from single individuals from a population stably maintaining *mptDf1* heteroplasmy (n=25). (E) mtDNA levels in individual L4 worms, normalized to actin and rank-ordered by *mptDf1* copy number. (F) Similar to *uaDf5*, wider variation in *mptDf1* relative to wildtype copy number (p<0.05) suggests that wildtype mtDNA, but not *mptDf1* mtDNA, is subject to homeostatic copy number control. AU, arbitrary units.

Here, we sought to distinguish between these possibilities of maintenance of total mtDNA copies versus wildtype mtDNA copies. Using ddPCR, we quantified mtDNA copy number from individual day 4 adults. mtDNA replication in *C. elegans* does not start until the L4 stage, after which it greatly increases during the first few days of adulthood and reaches steady state levels by day 4 (Fig. S2) (Bratic, Hench et al. 2009). mtDNA copy number measurements in day 4 adults reflect germline mtDNA since more than 95% of the mtDNA content in an adult hermaphrodite is in the germline syncytium, the shared cytoplasm of germ cells (Bratic, Hench et al. 2009). The ‘direct competition’ model predicts relatively constant levels of total mtDNA amongst individuals, with a trade-off between the amount of wildtype and *uaDf5* mtDNA copies. In contrast, our results show that wildtype mtDNA levels are remarkably consistent across individuals while *uaDf5* levels span a wide range. These findings are especially pronounced when animals are rank-ordered from lowest to highest *uaDf5* levels (Fig. 2B). Analysis of the range of wildtype and *uaDf5* mtDNA copy number shows substantially wider range in *uaDf5* mtDNA copy number compared to wildtype mtDNA (Fig. 2C). Taken together, our data show that *uaDf5* mtDNA levels increase in addition to, not at the expense of, wildtype mtDNA levels. These data also provide an explain for previously published data showing that total mtDNA levels in *uaDf5* heteroplasmic animals are higher than in homoplasmic wildtype animals (Tsang and Lemire 2002). In conclusion, our data are consistent with the hypothesis that wildtype mtDNA levels, but not the total mtDNA levels, are well regulated.

**Figure S2.**
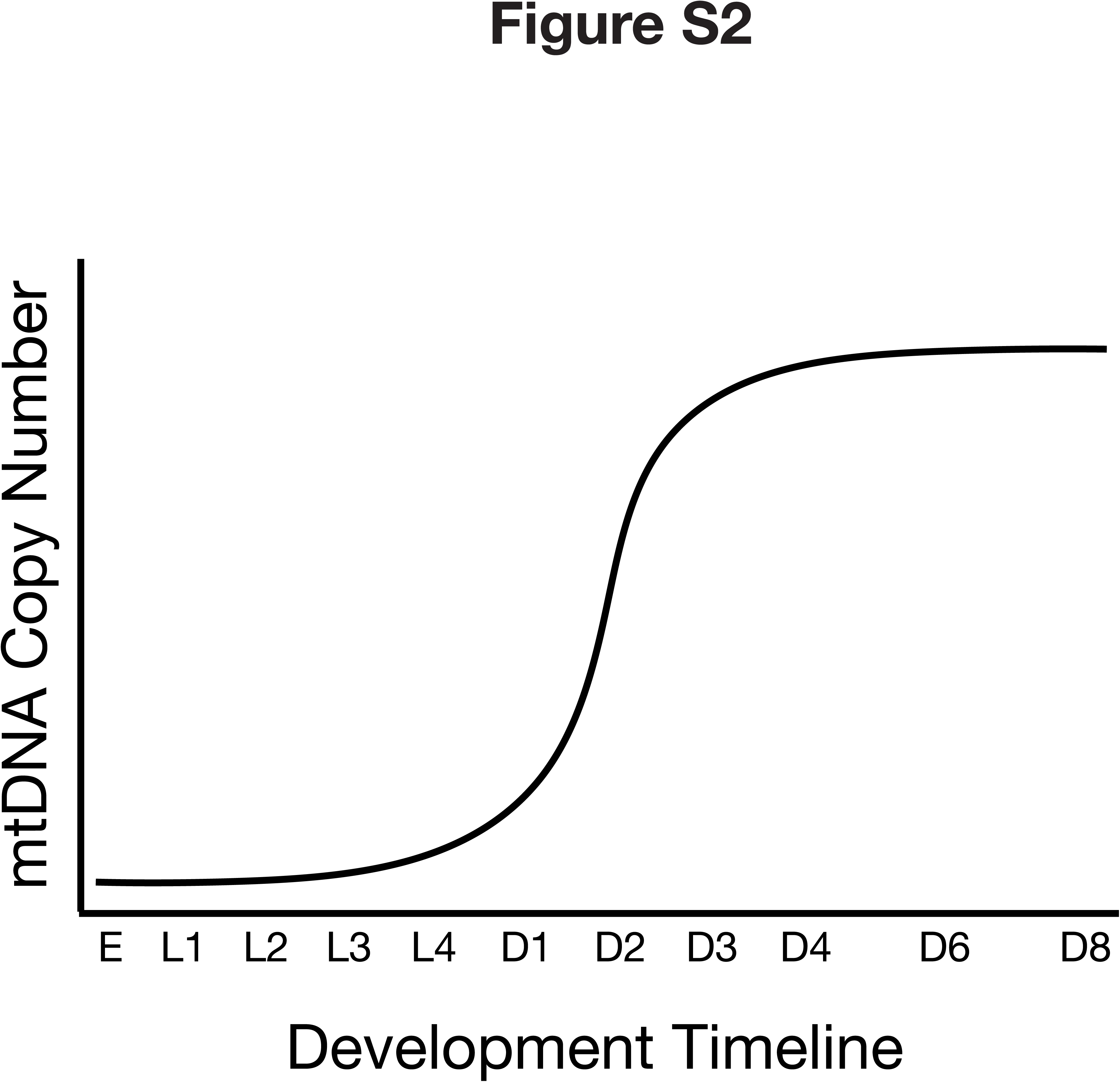
Schematic diagram of mtDNA levels as a function of development in*C. elegans*. Replication primarily occurs between the fourth larval stage (L4) and early adulthood (D1-D3), reaching steady state by day 4. Adapted from Bratic *et al.*, 2009.

### Small deletion bearing *mptDf1* mtDNA also has ‘runaway’ copy number dynamics

Experimental data from mammalian cell culture support the hypothesis that mutant mtDNA with large deletions like *uaDf5* might have a replicative advantage over wildtype mtDNA because of their smaller size (Wallace 1992, Moraes, Kenyon et al. 1999, Diaz, Bayona-Bafaluy et al. 2002). This advantage would allow the smaller genome to replicate faster, potentially giving rise to the ‘runaway’ dynamics that we observe for *uaDf5*. If this hypothesis were correct, we would expect to see different mtDNA dynamics in mtDNA harboring smaller deletions. To address this possibility, we investigated mtDNA copy number dynamics in animals that are heteroplasmic for *mptDf1*, a mutant mtDNA carrying a 179bp deletion (Fig. 1A), significantly smaller than the 3.1kb *uaDf5* deletion. We identified *mptDf1* deletion in a heteroplasmic strain from whole genome sequencing data of mutagenized worms from the Million Mutation Project (Thompson, Edgley et al. 2013). In contrast to the *uaDf5* deletion, which reduces mtDNA genome size by ~20%, the *mptDf1* deletion removes only ~1% of the genome. Interestingly, despite backcrossing to N2 reference strain five times, we noticed that high levels of *mptDf1* cause gonadal defects, and likely explain the lower average heteroplasmy frequency of *mptDf1* compared to *uaDf5* (Fig. S3). Despite this lower heteroplasmy frequency, and the fact that we had to use L4 stage individuals given germline defects in adults, we find that the mtDNA copy number dynamics in *uaDf5* and *mptDf1* heteroplasmies are remarkably similar, that is, wildtype mtDNA levels were relatively steady across individuals despite increasing levels of mutant mtDNA (Fig. 2D-F). While not ruling out replicative advantage, the data suggest that other mechanisms contribute to regulation of *uaDf5* and *mptDf1* copy number dynamics.

**Figure S3.**
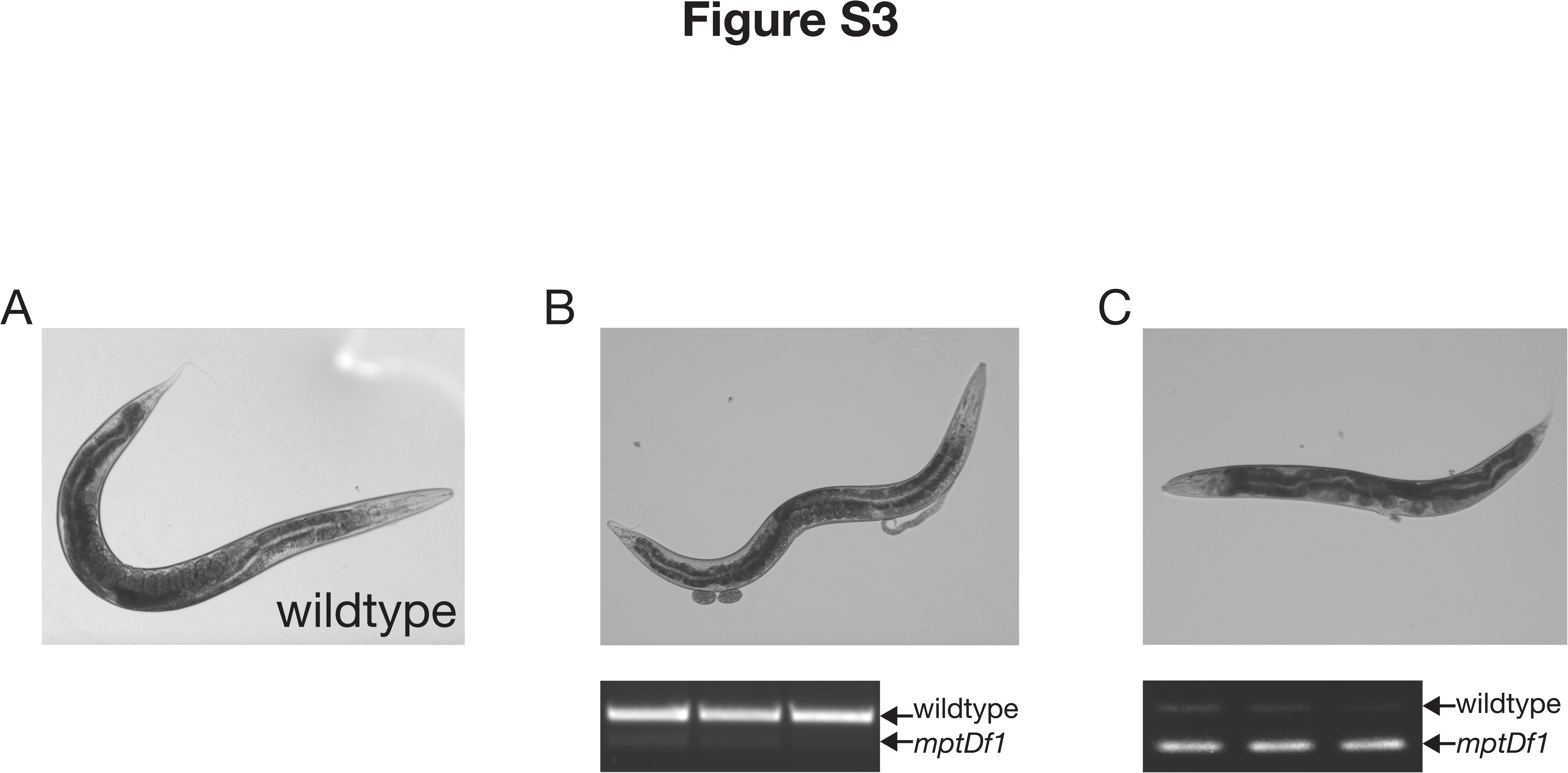
High*mptDf1* mtDNA levels affect gonad development in *C. elegans*. (A) Image of an adult animal homoplasmic for wildtype mtDNA (N2 strain). (B) Image of an adult animal heteroplasmic for low levels of *mptDf1* mtDNA. Three representative individuals with low *mptDf1* levels were lysed and their *mptDf1* levels assessed by PCR (gel image below). (C) Image of adult animal heteroplasmic for high levels of *mptDf1* mtDNA. Three representative individuals with high *mptDf1* levels were lysed and their *mptDf1* levels assessed by PCR (gel image below).

### mtDNA transcriptional imbalance in *uaDf5* heteroplasmic animals

Since heteroplasmic animals carry *uadf5* mtDNA in addition to wildtype mtDNA, we ascertained whether the *uaDf5* mtDNA contributed to the mtDNA transcripts. We reasoned that expression of all genes from the wildtype mtDNA but only some from the *uaDf5* genome would result in a stoichiometric imbalance in mtDNA-encoded transcript levels (see schematic Fig. 3A). We found that *CYTB* and *ND1*, genes deleted from *uaDf5*, were expressed at the same levels in homoplasmic wildtype animals as in *uaDf5* heteroplasmic animals (Fig. 3B). Transcript levels of the nuclear-encoded electron transport chain component *NUO2*, as well as actin, were also unaltered in the heteroplasmic animals (Fig. 3B). However, compared to homoplasmic wildtype animals, transcript levels of *COI*, *COII*, *COIII*, *ND4*, *ND5*, and *ND6*, which are still encoded by *uaDf5* mtDNA, were significantly elevated in the *uaDf5* heteroplasmic population (Fig. 3B). This implies that *uaDf5* mtDNA are transcriptionally active and their expression contributes to substantial transcriptional imbalances in mtDNA-encoded genes.

**Figure 3.**
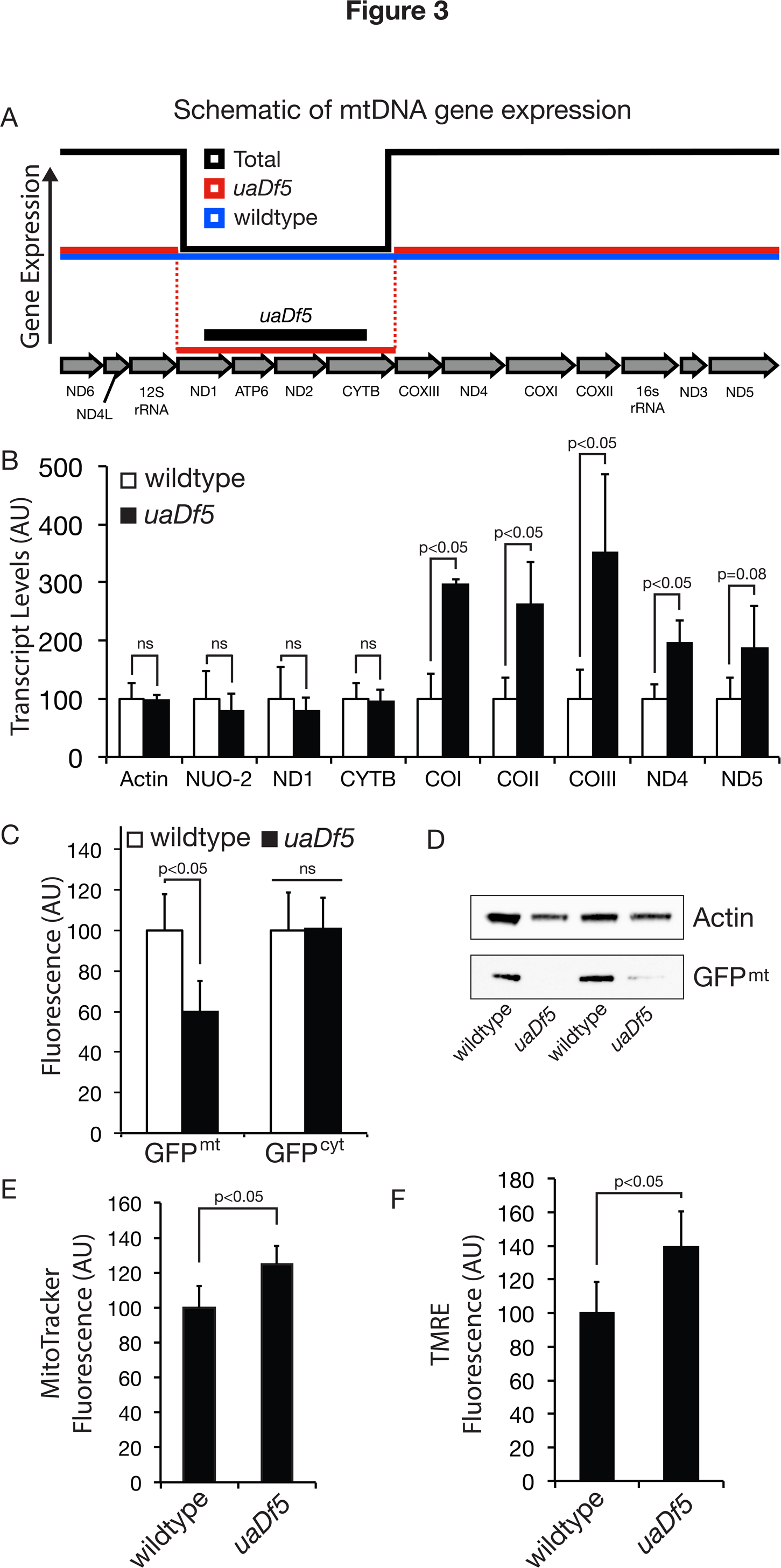
Mitochondrial function is perturbed in*uaDf5* animals. (A) Schematic showing expected expression of mtDNA-encoded transcripts. The presence of *uaDf5* mtDNA is expected to result in stoichiometric imbalance of gene expression, as the expression of *uaDf5* and wildtype mtDNA copies (red and blue lines, respectively) combine to generate total expression (black line) at elevated levels for genes located outside the deletion but at wildtype levels for genes missing from the *uaDf5* mtDNA. (B) Animals heteroplasmic for *uaDf5* exhibit expression levels similar to that of wildtype animals for mtDNA-encoded genes affected by the deletion (*CYTB* and *ND1*), as well as a nuclear-encoded mitochondrial gene (*NUO2*) and actin. However, *uaDf5* heteroplasmy results in overexpression for mtDNA-encoded genes located outside the *uaDf5* deletion (*COXI*, *COXII*, *COXIII*, *ND4*, and *ND5*). All transcript levels are normalized to wildtype. Error bars represent standard deviation. For each bar, n=4 lysates (pooled animals). (C) Mitochondrially targeted GFP (GFP^mt^), but not cytosolic GFP (GFP^cyt^), is significantly reduced in *uaDf5* heteroplasmic individuals (n=25). (D) Western blot analysis of wildtype and *uaDf5* heteroplasmic animals expressing GFP^mt^ reveals reduced levels in *uaDf5* heteroplasmic individuals relative to actin. Data are shown from two biological replicates each for wildtype and *uaDf5* strain. (E) Fluorescence increase in *uaDf5* animals stained with mitochondrial membrane potential independent dye MitoTracker Green FM (n=21) and (F) membrane potential dependent dye TMRE (n=25). Error bars represent standard deviation. AU, arbitrary units.

### Mitochondrial perturbations in *uaDf5* heteroplasmic animals

*uaDf5* affects organismal fitness; *uaDf5* animals have decreased egg-laying and defecation rates, reduced lifespan, and decreased sperm motility (Liau, Gonzalez-Serricchio et al. 2007). At the molecular level, we observe overexpression of *uaDf5* mtDNA-encoded transcripts, resulting in transcriptional imbalance. Given these effects, we sought to determine whether *uaDf5* has cellular consequences. Mitochondrially targeted green fluorescence protein (GFP^mt^) has previously been used as a model matrix protein to assess mitochondrial proteostasis (Yoneda, Benedetti et al. 2004, Benedetti, Haynes et al. 2006). We reasoned that if *uaDf5* animals have altered proteostasis, it might result in decreased fluorescence of GFP^mt^. Indeed, we observe a significant decrease in fluorescence in *uaDf5* animals that express GFP^mt^ in the intestinal cells under the control of the *ges-1* promoter (Fig. 3C) (Benedetti, Haynes et al. 2006). No such decrease is observed in fluorescence of cytoplasmic GFP (GFP^cyt^) in *uaDf5* animals (Fig. 3C). These data suggest that the fluorescence of mitochondrially targeted GFP is specifically affected in *uaDf5* animals. Consistent with fluorescence data, western blot analysis shows decreased GFP^mt^ protein levels in *uaDf5* animals (Fig. 3D). In contrast to decreased GFP^mt^, we observe increased staining in *uaDf5* animals with fluorescent dye MitoTracker Green FM, which localizes to mitochondria independent of the membrane potential (Hicks, Howe et al. 2012, Dingley, Polyak et al. 2014), as well as with tetramethyl rhodamine ethyl ester (TMRE) (Yoneda, Benedetti et al. 2004, Billing, Kao et al. 2011, Dingley, Polyak et al. 2014, Palikaras, Lionaki et al. 2015), which localizes to mitochondria in a membrane potential dependent manner (Fig. 3E-F). These data suggest an increase in mitochondrial organelle mass in *uaDf5* animals, correlating with an increase in total mtDNA levels. These data also suggest that the decreased GFP^mt^ levels and fluorescence in *uaDf5* animals are due to alterations in mitochondrial proteostasis rather than due to decreased mitochondrial organelle mass. This altered proteostasis might reflect decreased GFP^mt^ import efficiency into mitochondria or a compromised protein-folding environment inside the mitochondrial matrix. Either scenario might result in degradation of the unfolded/misfolded GFP^mt^. Taken together, these data suggest mitochondrial alterations in *uaDf5* animals.

### High levels of *uaDf5* activate the mitochondrial unfolded protein response

The mitochondrial unfolded protein response (UPR^mt^) has emerged as an important protective stress response that is activated under a variety of conditions that affect mitochondrial proteostasis (Yoneda, Benedetti et al. 2004, Haynes and Ron 2010, Baker, Nargund et al. 2012, Houtkooper, Mouchiroud et al. 2013, Runkel, Liu et al. 2013). This homeostatic response involves expression of hundreds of target genes including chaperones and proteases that eliminate misfolded and nonfunctional complexes inside mitochondria (Nargund, Pellegrino et al. 2012, Nargund, Fiorese et al. 2015). To determine whether UPR^mt^ is induced in *uaDf5* animals, we quantified *hsp-6* and *hsp-60* transcript levels in *uaDf5* animals. HSP-6 and HSP-60 are mitochondrial chaperones that are induced by UPR^mt^ (Yoneda, Benedetti et al. 2004, Nargund, Pellegrino et al. 2012). We observed significant increase in *hsp-6* and *hsp-60* transcript levels in *uaDf5* animals (Fig. 4A). We were able to confirm this induction with a transgenic fluorescence reporter in which the *hsp-60* promoter drives GFP expression, although UPR^mt^ induction is variable across individuals (Fig. 4B-C) (Yoneda, Benedetti et al. 2004). These data suggest that UPR^mt^ might be appreciably induced in animals harboring *uaDf5* levels above a certain threshold. Consistent with this notion, we were able to observe a weak but positive relationship between UPR^mt^ activation and *uaDf5* levels (Fig. 4D). The lack of a stronger correlation might be a product of the fact that most of the animals we analyzed have *uaDf5* levels within a very narrow range of 60-80%. Taken together, these results suggest that presence of the selfish mtDNA *uaDf5* induces UPR^mt^.

**Figure 4.**
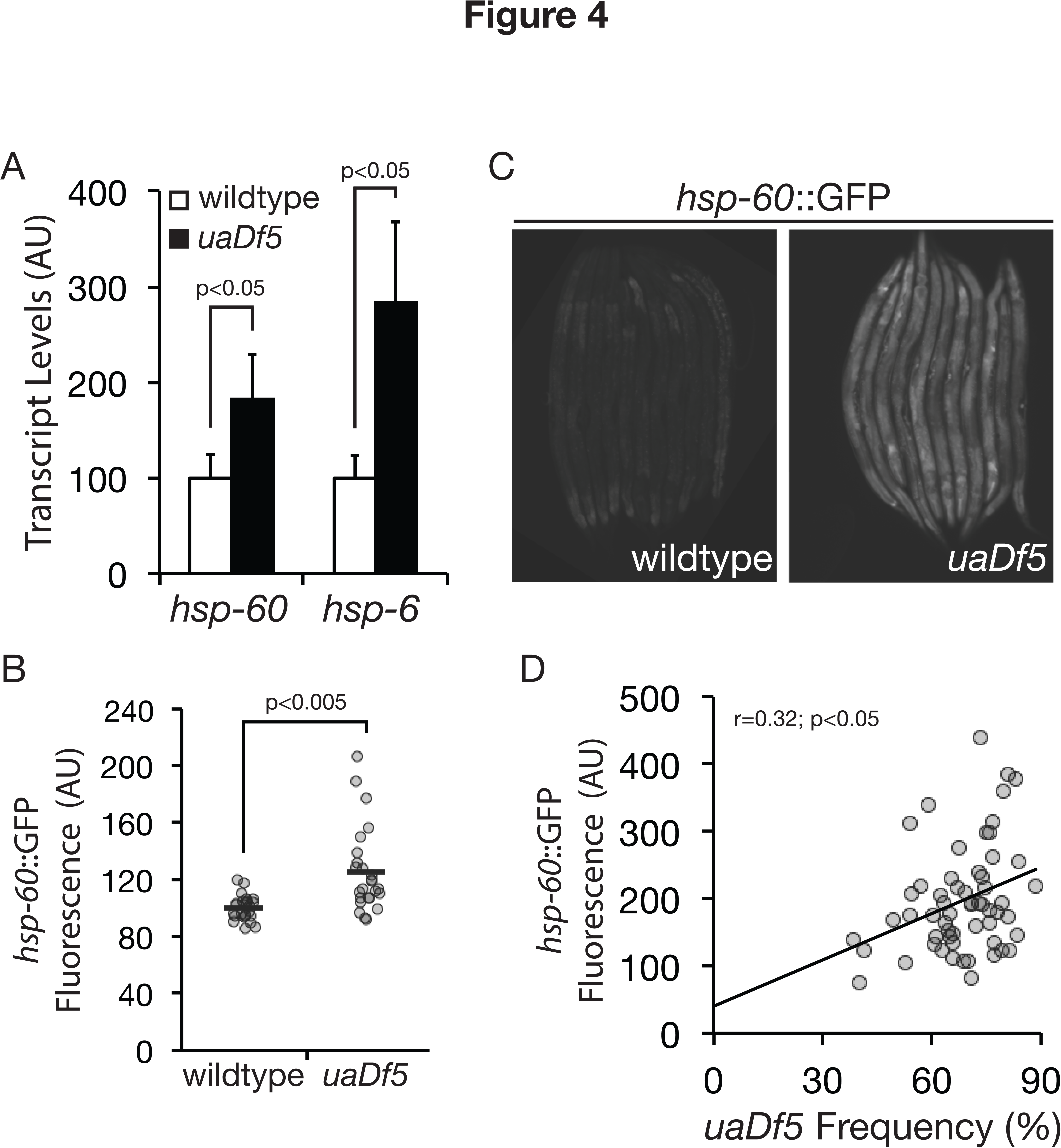
UP^mt^ is activated in heteroplasmic animals carrying *uaDf5* mtDNA. (A) Transcription of two UPR^mt^-activated molecular chaperones (*hsp-60* and *hsp-6*) is increased in individuals with *uaDf5* compared to wildtype individuals. For each bar, n=4 lysates (pooled animals). (B) Quantification of fluorescence between wildtype homoplasmic and *uaDf5* heteroplasmic animals shows increased activation of the UPR^mt^ marker *hsp-60*:GFP in the presence of *uaDf5* mtDNA (n=25). Each data point is from a single individual picked randomly from a population. (C) Visual comparison of GFP fluorescence between *uaDf5* and wildtype animals, each expressing *hsp-60*::GFP. Wildtype animals were picked at random from a population but only *uaDf5* animals with apparent fluorescence were picked to show UPR^mt^ activation. (D) Positive relationship between *uaDf5* frequency and *hsp-60*::GFP fluorescence (trendline) indicates that UPR^mt^ activation increases at higher *uaDf5* frequency. Each data point corresponds to a single individual (n=58). Error bars represent standard deviation. AU, arbitrary units.

### UPR^mt^ modulates *uaDf5* heteroplasmy levels

UPR^mt^ plays a protective role under conditions that affect mitochondria (Baker, Nargund et al. 2012, Nargund, Pellegrino et al. 2012, Runkel, Liu et al. 2013). Might the protective role for UPR^mt^ inadvertently create the conditions that allow *uaDf5* to achieve high levels in certain individuals? According to this hypothesis, we predict that loss of UPR^mt^ would result in a decrease in *uaDf5* levels. To test this hypothesis, we conducted RNAi-mediated knockdown of *atfs-1*, a gene that encodes a transcription factor central to UPR^mt^ induction (Nargund, Pellegrino et al. 2012). We observed a significant decrease in the average *uaDf5* frequency in *atfs-1* knockdown animals (Fig. 5A). We similarly observe a decrease in *uaDf5* levels in *atfs-1* loss-of-function mutants (Fig. 5B).

**Figure 5.**
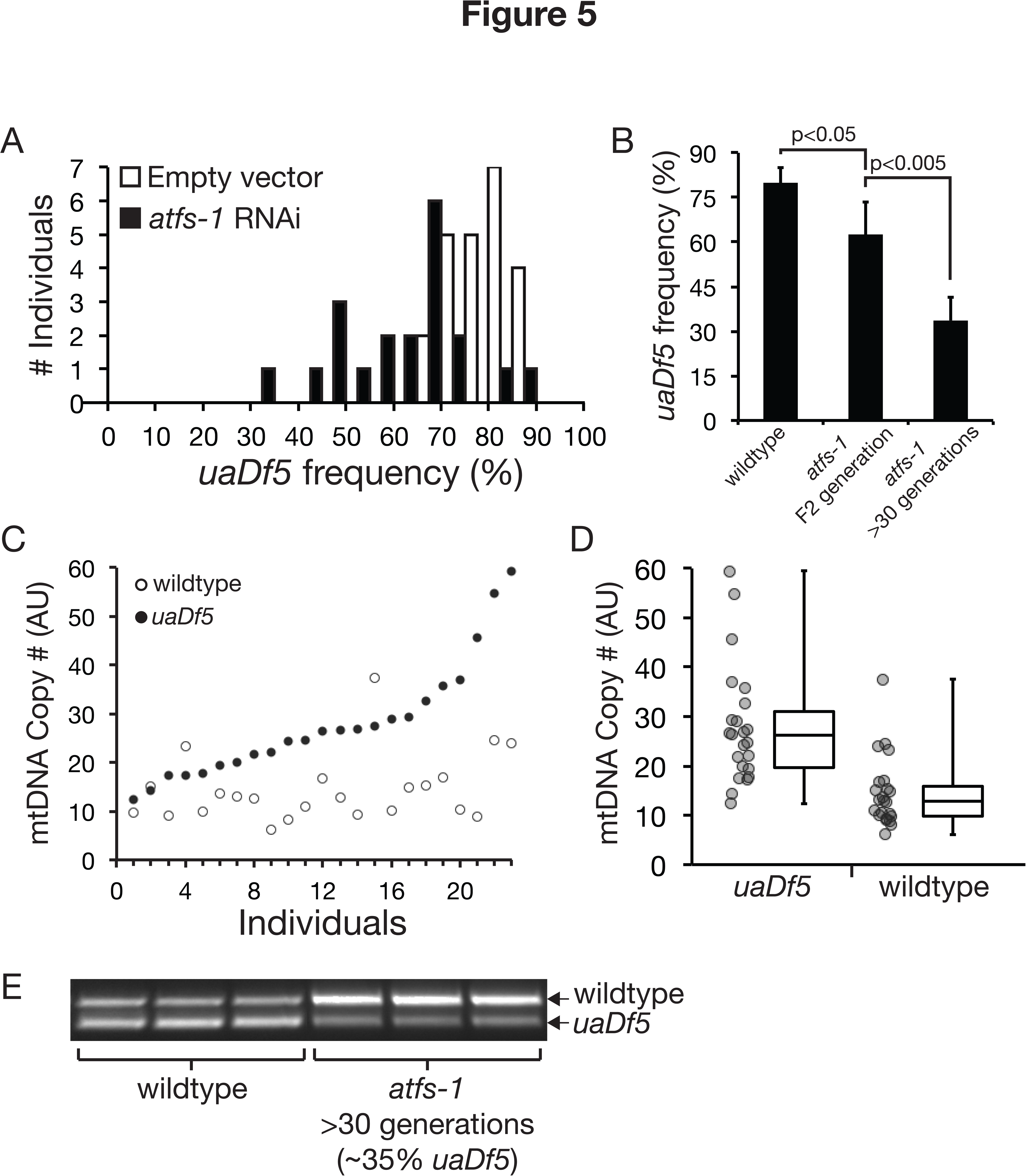
Loss of UP^mt^ activation results in decreased *uaDf5* levels but does not affect mtDNA copy number control. (A) Growth under RNAi-mediated knockdown of *atfs-1* (n=20), required for UPR^mt^ activation, results in a shift to lower *uaDf5* frequency relative to growth under control conditions (n=23; p<0.05). (B) *uaDf5* frequency decreases in heteroplasmic animals homozygous for the *atfs-1*(*tm4525*) loss-of-function allele (n=16) compared to heteroplasmic animals that express wildtype *atfs-1* (n=16), in which high *uaDf5* levels are stably maintained. *uaDf5* frequency decreases further in the *atfs-1* null animals after multiple generations (n=4 lysates, pooled animals) but is not lost completely. (C) Quantification of mtDNA copy number in individual day 4 adult animals homozygous for the *atfs-1* loss-of-function allele (n=23), normalized to actin and rank-ordered by *uaDf5* mtDNA copy number. (D) Wider variation in *uaDf5* relative to wildtype copy number (p<0.05) in *atfs-1* null animals from panel C suggests that mtDNA copy number control persists in absence of UPR^mt^. (E) PCR of single heteroplasmic individuals against the *atfs-1* wildtype or *atfs-1* null nuclear background shows that *uaDf5* is retained in both lines but is at lower levels in the null animals after about 30 generations. Note that because mutant and wildtype templates compete for amplification, the wildtype band appears fainter when *uaDf5* levels are high but does not actually reflect reduced wildtype mtDNA levels (see Fig. 2B). Error bars represent standard deviation. AU, arbitrary units.

Quantification of mtDNA dynamics shows that wildtype mtDNA levels are well regulated in *atfs-1* loss-of-function mutants, thus ruling out loss of mtDNA copy number control as a potential explanation for the decrease in *uaDf5* levels (Fig. 5C-D). Given that UPR^mt^ is likely induced in animals harboring *uaDf5* levels above a certain threshold, and the hypothesis that *uaDf5* might exploit additional mechanisms to persist, such as mtDNA copy number control, we predict that although *uaDf5* levels decrease in absence of UPR^mt^, *uaDf5* would still be able to persist. Indeed, *uaDf5* is still present, albeit at low levels, in *atfs-1* loss-of-function animals that we have continuously maintained for more than 30 generations (Fig. 5E). These data are consistent with the hypothesis that loss of UPR^mt^ exposes *uaDf5* to more stringent selection, which causes a decrease in *uaDf5* levels but not its complete elimination.

If UPR^mt^ activation allows for tolerance to high levels of *uaDf5*, then we predict that restoring *atfs-1* will allow *uaDf5* levels to recover after *atfs-1* RNAi treatment. Indeed, *uaDf5* levels recover within a single generation when *atfs-1* expression is restored in *uaDf5* animals after growing for several generations on *atfs-1* RNAi conditions (Fig. 6A). These data suggest that UPR^mt^ is required for *uaDf5* to attain high levels. We next tested if the converse was also true, *i.e.*, whether constitutive activation of UPR^mt^ decreases selection against *uaDf5*, thereby driving *uaDf5* to higher frequency. For this, we tested animals heterozygous for an *atfs-1* gain-of-function allele to constitutively activate UPR^mt^ (Rauthan, Ranji et al. 2013). We did not observe any significant increase in the *uaDf5* levels in animals heterozygous for the *atfs-1* gain-of-function allele (Fig. 6B). However, given that the starting *uaDf5* levels in our *uaDf5* strain are already very high (approximately 80%), we speculated that they might not be able to increase substantially due to an upper threshold effect. It is also be possible that given the induction of UPR^mt^ in individuals with high levels of *uaDf5*, the *atfs-1* gain-of-function allele might not induce significant further UPR^mt^ activation in these animals. To overcome these limitations, we tested whether *uaDf5* levels can rise in *atfs-1* gain-of-function heterozygotes in a population with lower starting *uaDf5* frequency (approximately 30%). In this case, we observed a significant increase in *uaDf5* frequency in *atfs-1* gain-of-function heterozygotes relative to animals homozygous for wildtype *atfs-1* (Fig. 6C). Together, our results show that *atfs-1* can modulate *uaDf5* levels.

**Figure 6.**
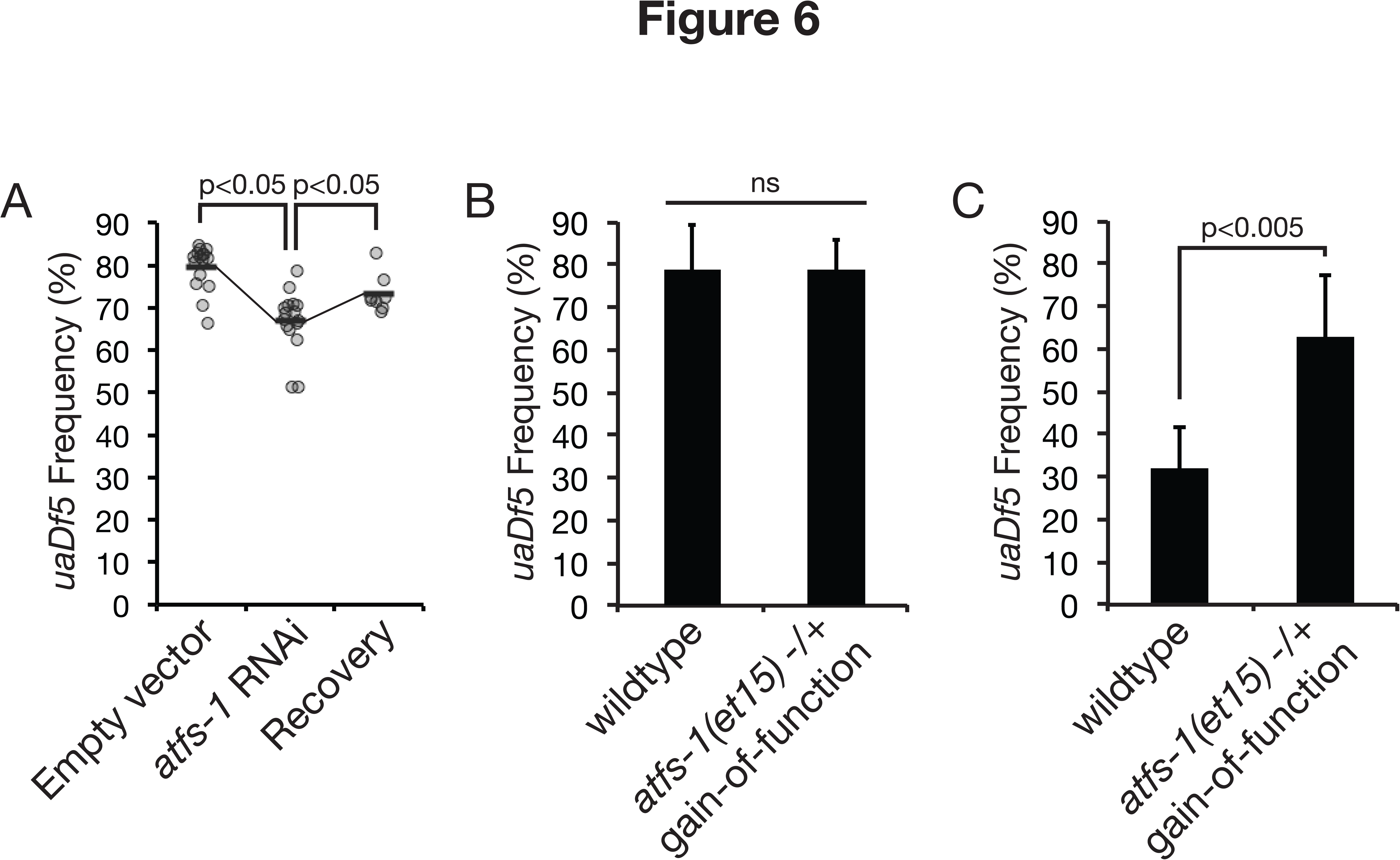
Persistence of*uaDf5* at high frequency depends in part on UPR^mt^ activation. (A) Growth under RNAi-mediated knockdown of *atfs-1* across seven generations reduces average *uaDf5* frequency (n=16). However, restoration of *atfs-1* expression by returning *atfs-1* knockdown animals to control conditions results in recovery of elevated *uaDf5* frequency in a single generation (n=8 lysates, pooled animals). (B) When starting *uaDf5* frequency is high (75-80%), constitutive UPR^mt^ activation in individuals heterozygous for an *atfs-1* gain-of-function allele (n=12) causes no further rise in average *uaDf5* frequency relative to the wildtype background (n=12); (C) however, *uaDf5* frequency rises when the *atfs-1* gain-of-function allele is crossed into a strain harboring lower *uaDf5* levels (~30%; n=8). Error bars represent standard deviation.

### Loss of UPR^mt^ does not select against *uaDf5* at the organismal level

Increased sensitivity of animals with high *uaDf5* levels to *atfs-1* loss at the organismal level provides one potential explanation for the observed decrease in *uaDf5* levels. RNAi knockdown of *atfs-1* enhances developmental delay in animals exposed to paraquat (Runkel, Liu et al. 2013). Mitochondrial-stressed *isp-1(qm150)* and *clk-1(qm30)* mutants similarly fail to develop under *atfs-1* RNAi conditions (Nargund, Pellegrino et al. 2012). We sought to determine whether knockdown of *atfs-1* RNAi causes similar developmental delay in *uaDf5* animals. While we observe a mild developmental delay in *uaDf5* animals, it is not enhanced by *atfs-1* knockdown (Fig. 7A). We also did not observe appreciable levels of embryonic lethality in *uaDf5* animals raised on *atfs-1* RNAi (Fig. 7B). Finally, there is no increase in lethality up to day 4 of adulthood (Fig. 7C), which is when we assess and observe a decrease in *uaDf5* levels on *atfs-1* RNAi. The absence of an *atfs-1* knockdown-dependent effect on reproduction suggests that selection against high *uaDf5* levels is unlikely to operate at the organismal level.

**Figure 7.**
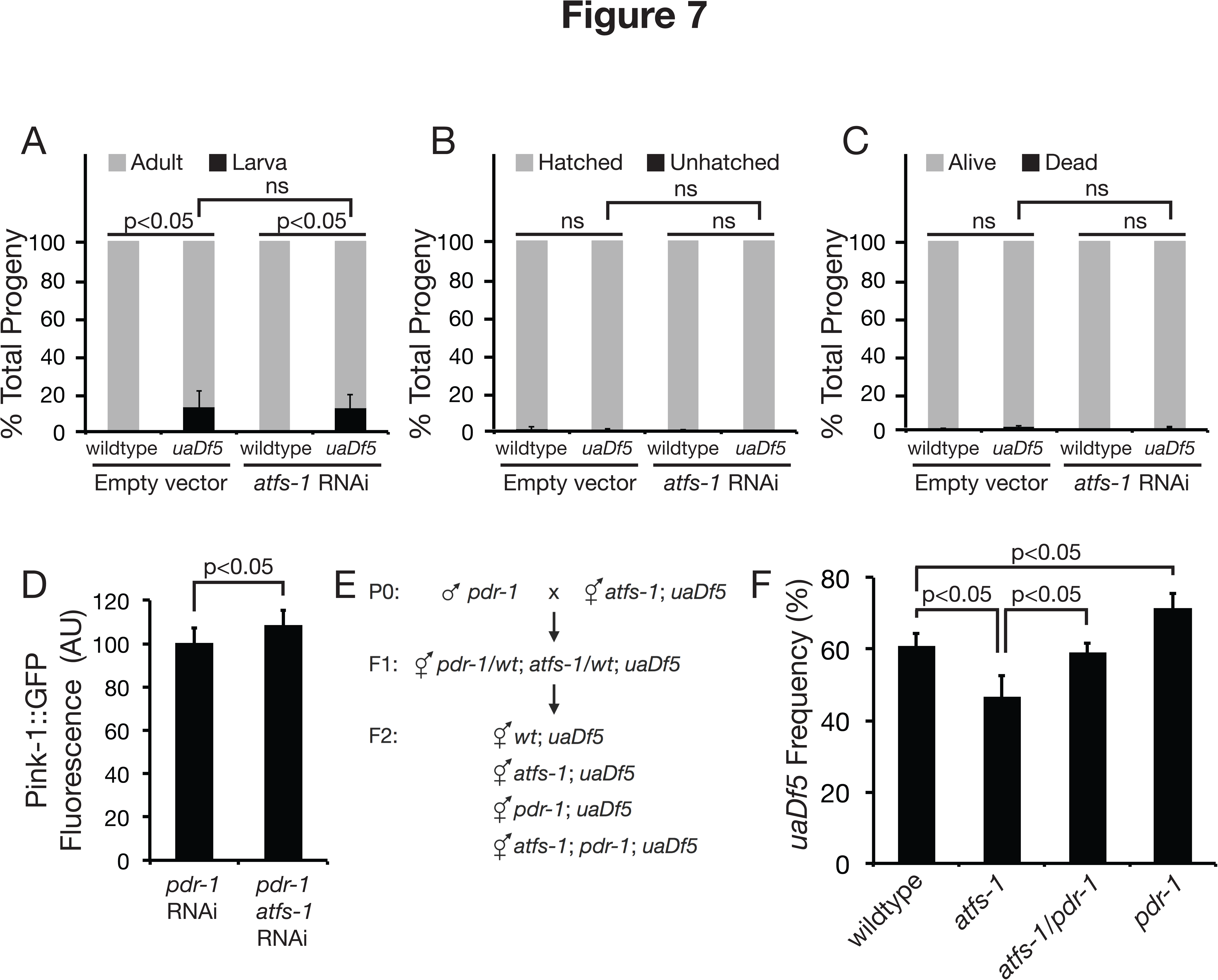
UP^mt^ protects *uaDf5* from mitophagy. (A) Heteroplasmic individuals exhibit delayed growth as 100% of progeny from wildtype parents reach adulthood in three days, approximately 10% of *uaDf5* progeny remain in the larval stage. Knockdown of *atfs-1* showed no effect on development in homoplasmic wildtype animals and did not further enhance developmental delay in *uaDf5* heteroplasmic animals. (B) No significant difference was observed between *uaDf5* and wildtype animals, or between *atfs-1* knockdown and control conditions, on the percentage of embryos that remain unhatched after one day or (C) on the percentage of lethality among day 4 adults. (D) Quantification of Pink-1::GFP fluorescence shows increased mitophagy in *uaDf5* animals upon *pdr-1;atfs-1* double knockdown (n=10) compared to knockdown of *pdr-1* alone (n=10). AU, arbitrary units. (E) Crossing scheme employed to isolate *uaDf5* animals in wildtype, *atfs-1* null, *pdr-1* null, and *atfs-1;pdr-1* double mutant backgrounds. (F) Quantification of *uaDf5* levels shows recovery of *uaDf5* levels in *atfs-1;pdr-1* double mutants compared to *atfs-1* single mutant animals. *uaDf5* recovers to the highest levels in *pdr-1* single mutants. For each bar, n=4 lysates (pooled animals). Error bars represent standard deviation. AU, arbitrary units.

### UPR^mt^ modulates *uaDf5* levels via mitophagy

Organelle level selection provides an alternate possibility for the observed decrease in *uaDf5* levels in absence of *atfs-1*. According to this hypothesis, *uaDf5* might be more susceptible to mitophagy in absence of UPR^mt^. Mitophagy is initiated by stabilization of Pink-1 on dysfunctional mitochondria, which in turn recruits parkin to mediate mitophagy (Randow and Youle 2014). Here, we observe an increase in Pink-1::GFP fluorescence in *uaDf5* animals with RNAi knockdown of *atfs-1* compared to control (Fig. 7D; these experiments were conducted in parkin homolog *pdr-1* RNAi background to stabilize Pink-1::GFP localization to mitochondria). The increase in Pink-1::GFP fluorescence suggests that loss of UPR^mt^ promotes mitophagy in *uaDf5* animals. If mitophagy mediates decrease in *uaDf5* levels in absence of UPR^mt^, then we predict that *uaDf5* levels should recover in animals without UPR^mt^ that also have compromised mitophagy. In order to test this prediction, we quantified *uaDf5* levels in *atfs-1; pdr-1* double mutants (Fig. 7E-F). We observed significant recovery of *uaDf5* levels in *atfs-1; pdr-1* double mutant animals compared to *atfs-1* single mutants. Interestingly, we saw the greatest recovery of *uaDf5* levels in *pdr-1* single mutants (Fig. 7F). These data are consistent with recently published observation that *uaDf5* frequency increases in *pdr-1* mutants (Valenci, Yonai et al. 2015). In summary, our genetic analyses indicate that UPR^mt^ protects *uaDf5* from mitophagy, with loss of UPR^mt^ exposing it to mitophagy.

## Discussion

Mechanisms that regulate selfish mutant mtDNA dynamics are poorly understood. Here, we address this issue by establishing that an mtDNA mutation in *C. elegans*, *uaDf5*, behaves as a selfish genetic element. We show that the ‘runaway’ copy number dynamics of *uaDf5* are consistent with it exploiting the host’s homeostatic mtDNA copy number control mechanism to achieve high frequency. We further show that the UPR^mt^, a mitochondrial stress response system, is activated in animals with high *uaDf5* levels. *uaDf5* frequency decreases in absence of UPR^mt^, suggesting that the *uaDf5* mutant also exploits UPR^mt^ to persist at high levels. Thus, *uaDf5* can be viewed as exploiting the very homeostatic responses that its presence activates to persist and proliferate (Fig. S4). Such exploitation of endogenous pathways provides a potential explanation for how mutant mtDNA are able to achieve high frequency to cause disease in humans.

**Figure S4.**
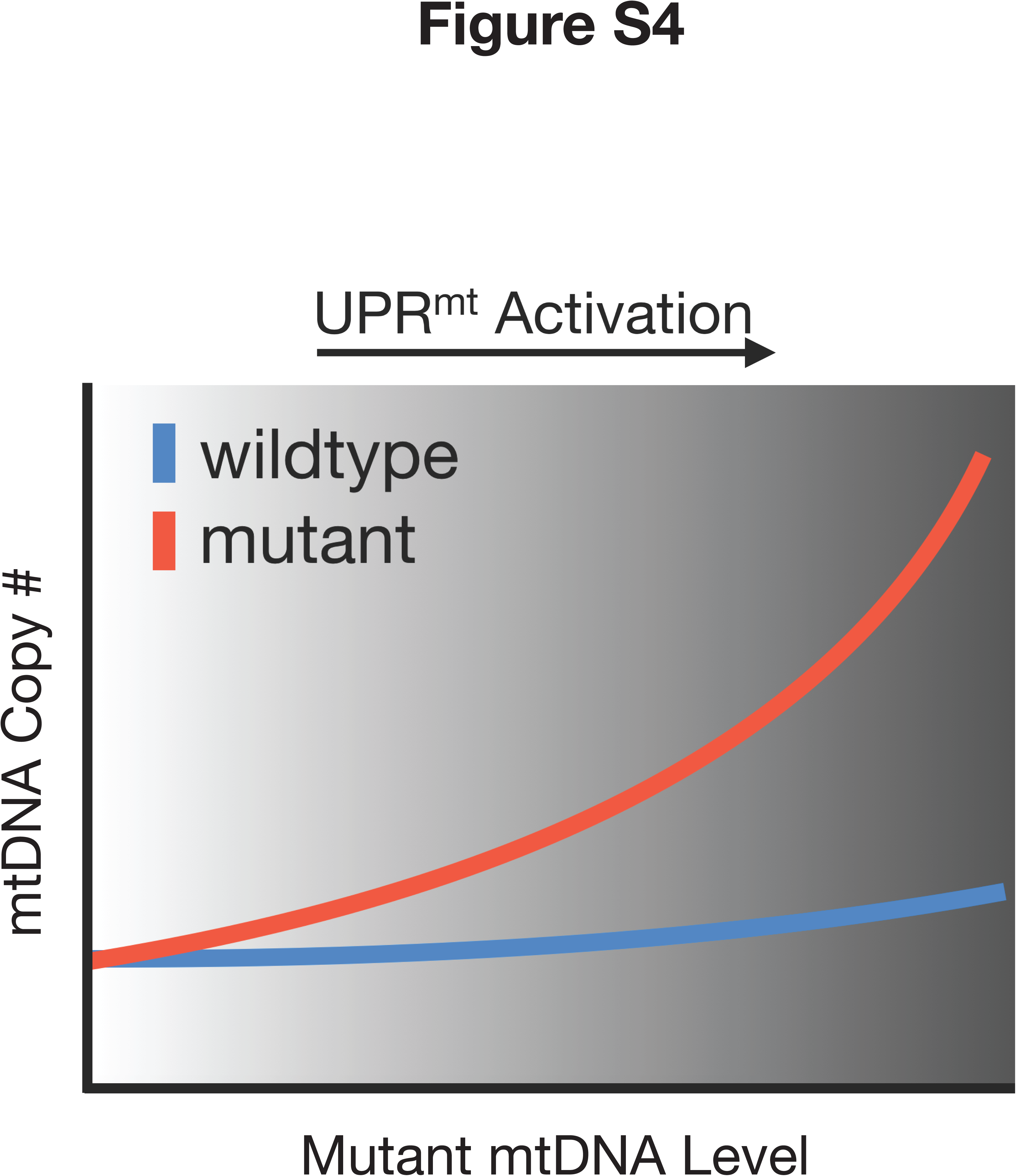
Proposed model illustrating the heteroplasmy dynamics of a selfish mtDNA mutant. Selfish mutant mtDNA exploit mtDNA copy number control to ‘hitchhike’ to high frequency, as wildtype mtDNA levels remain relatively constant. UPR^mt^ allows mutant mtDNA to persist at high levels.

### mtDNA copy number control

Our investigation of mtDNA copy number dynamics in a *uaDf5* heteroplasmic *C. elegans* strain shows regulation of wildtype mtDNA levels but ‘runaway’ dynamics of *uaDf5*. What mechanisms can explain these observations? We find that our data are most consistent with a previously proposed Maintenance of Wild Type model (Chinnery and Samuels 1999, Capps, Samuels et al. 2003, Kowald and Kirkwood 2014). This model requires a feedback mechanism whereby mtDNA replication rate is inversely related to wildtype mtDNA levels (Fig. S5). This feedback mechanism allows cells to maintain wildtype mtDNA levels at a set target. When mutant mtDNA fail to support production of this output, insufficient levels of wildtype mtDNA in a heteroplasmy induce mtDNA replication. However, random sampling by the replication machinery can lead to replication of mutant mtDNA, requiring additional rounds of replication and, in turn, production of more mutant mtDNA. Thus, this negative feedback model of copy number homeostasis predicts ‘runaway’ dynamics of mutant mtDNA, exactly like those observed for *uaDf5*. Based on these data, *uaDf5* can be viewed as taking advantage of homeostatic mtDNA copy number control to hitchhike to high frequency. Similar over-proliferation of mtDNA is observed in individuals with heteroplasmic mtDNA diseases and gives muscles a ‘ragged red fiber’ appearance (Durham, Samuels et al. 2007). Taken together, these data suggest that exploitation of homeostatic mtDNA copy number control by mutant mtDNA might be a conserved and widespread strategy. Ours is the first report to show that this strategy can operate in the germline to regulate heteroplasmy transmission dynamics.

**Figure S5.**
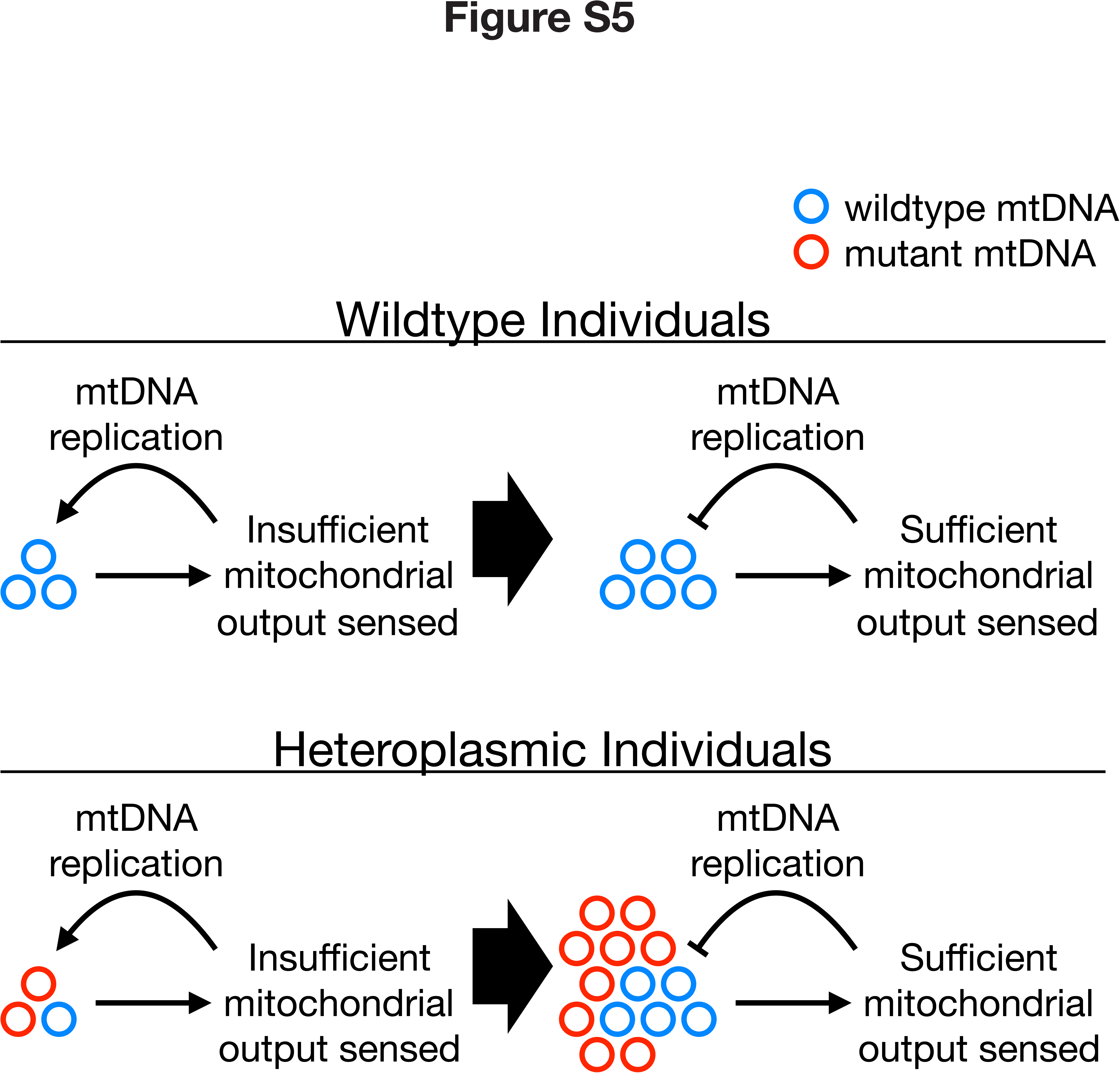
‘Maintenance of wild type’ model of homeostatic mtDNA copy number regulation. In wildtype individuals (top), an output of mtDNA is sensed, triggering feedback inhibition of replication when mtDNA copy number reaches a minimum threshold. In heteroplasmic individuals (bottom), mutant mtDNA copies that fail to contribute the same output are invisible to the mechanism of feedback inhibition. Consequently, the mutant copies hitchhike to higher frequency as the cell undergoes further mtDNA replication until sufficient wildtype levels are achieved. This model predicts that wildtype mtDNA copy number homeostasis is achieved at the cost of rising mutant levels.

We observed similar mtDNA dynamics in *mptDf1* heteroplasmic animals, which harbors a very small deletion compared to *uaDf5*. These data suggest that this mutant mtDNA also exploits mtDNA copy number control. While these data suggest that it is not necessary to invoke replicative advantage to explain *uaDf5* and *mptDf1* dynamics, our data do not rule out a role for such mechanisms in contributing to persistence of selfish mtDNA. Another model that ascribes an inherent advantage to mutant mtDNA was recently proposed (Kowald and Kirkwood 2014). According to this model, the production of the output sensed to ‘count’ mtDNA might be coupled to inhibition of replication of that specific mtDNA molecule (Kowald and Kirkwood 2014). This feature would have the effect of hastening runaway dynamics of mutant mtDNA. Our data does not exclude this possibility and it would be interesting to investigate the role of such coupling in the future.

What is the mtDNA output that is sensed to achieve mtDNA copy number control? Given that ND1 is the only gene deleted from both *uaDf5* and *mptDf1* mtDNA, it is possible that some output that is dependent on ND1 function is required to ‘count’ mtDNA. However, ND1 is one subunit of the large electron transport chain complex I. Hence, it is also possible that the output is produced by this complex and mutations in any one of the mtDNA encoded complex I subunits will result in the same runaway copy number dynamics. Consistent with a role of complex I, analysis of mtDNA deletions in mammals implicates components of complex I in regulating mtDNA copy number (Kowald and Kirkwood 2014). Finally, it is also possible that production of the sensed output is dependent on the entire electron transport chain. In this instance, mutations in any of the mtDNA-encoded genes will result in failure of the mutant mtDNA to contribute to the output. Future experiments with additional heteroplasmies with mutations or deletions in different mtDNA-encoded genes will shed light on this issue. Failure to contribute to the sensed output used for mtDNA copy number control might be a common feature of selfish mtDNA.

mtDNA copy number control has to occur during periods of mtDNA replication. Previous work has shown that mtDNA replication starts in L4 stage and continues throughout adulthood (Bratic, Hench et al. 2009). This replication coincides with germline proliferation. Furthermore, there is almost no increase in mtDNA copy number during these stages of development in germline free animals, suggesting that almost all mtDNA replication occurs in the developing germline (Bratic, Hench et al. 2009). Based on this data, we suggest that mtDNA copy number control occurs in the germline syncytium either continuously from L4 stage onwards, or during adulthood after mtDNA has reached steady state levels.

It is interesting to note a slight increase in wildtype mtDNA levels with increasing *uaDf5* levels. These data might be reflective of overcompensation in wildtype mtDNA replication. For instance, interference with wildtype mtDNA function by *uaDf5*-encoded products might result in an effective decrease in the number of functional wildtype mitocdhondria. Such interference might result in a cellular effort to increase levels of wildtype mtDNA above baseline.

### UPR^mt^ in *uaDf5* heteroplasmy

We observed UPR^mt^ activation in animals with high *uaDf5* levels as determined using a transgenic reporter of the *hsp-60* promoter driving GFP expression. We confirmed this UPR^mt^ activation by measuring *hsp-6* and *hsp-60* transcript levels. How does *uaDf5* induce UPR^mt^? Defective protein handling inside mitochondria is believed to trigger UPR^mt^ (Yoneda, Benedetti et al. 2004, Houtkooper, Mouchiroud et al. 2013, Mouchiroud, Houtkooper et al. 2013). Here, we observe an overexpression of mtDNA-encoded transcripts that are still intact in *uaDf5* mtDNA. If these transcripts are translated, they might overwhelm the protein-handling environment in the mitochondria.

Alternatively, they might result in formation of stoichiometrically imbalanced nonfunctional electron transport chain complexes. Decrease in GFP^mt^ levels and fluorescence in *uaDf5* animals is consistent with the hypothesis that either import of GFP^mt^ into mitochondria or its folding inside the matrix are compromised. Future experiments aimed at characterizing the details of protein-handling environment in *uaDf5* animals will help determine the mechanistic basis of UPR^mt^ induction.

UPR^mt^ activation relies on decreased import efficiency of ATFS-1 into mitochondria. Active protein import into mitochondria is known to rely on the mitochondrial membrane potential. In *uaDf5* animals, although we see UPR^mt^ induction and a role for ATFS-1 in modulating *uaDf5* heteroplasmy levels, mitochondrial membrane potential is likely not compromised. Indeed, we actually see an increase in staining of *uaDf5* animals with TMRE, a mitochondrial membrane potential dependent dye. This increase might not be reflective on an actual increase in mitochondrial membrane potential but probably results from an overall increase in mitochondrial organelle mass since we also observe an increase in staining with MitoTracker Green FM, a dye that localizes to mitochondria independent of the membrane potential. Overall, these data suggest a different molecular basis for decreased ATFS-1 import efficiency than decreased mitochondrial membrane potential. This conclusion is perhaps not surprising given that ATFS-1 import efficiency has not been linked to decreased membrane potential (Nargund, Pellegrino et al. 2012).

Our data show that UPR^mt^ is an important regulator of *uaDf5* levels. Why does loss of UPR^mt^ result in reduced *uaDf5* levels? We suggest that UPR^mt^ protects *uaDf5* against selection, with loss of UPR^mt^ sensitizing the background and exposing *uaDf5* to selection. Selection can operate at the organismal level, which would result in elimination of animals with high *uaDf5* levels either through developmental delay or lethality. While we observe a slight developmental delay in *uaDf5* animals, this delay is not enhanced by *atfs-1* RNAi. We also did not observe enhancement in embryonic lethality or increased death up till day 4 of adulthood when we assessed decreased *uaDf5* levels in *atfs-1* RNAi. These data suggest that UPR^mt^ ‘protects’ *uaDf5* against selection, perhaps at the organelle level. Under this hypothesis, mitochondria harboring high levels of *uaDf5* might be more susceptible to mitophagy in absence of UPR^mt^. In support of this hypothesis, we observe an increase in Pink-1::GFP fluorescence in *uaDf5* animals without the ability to induce UPR^mt^. We also observed recovery of *uaDf5* levels in *atfs-1; pdr-1* double mutants that lack the ability to induce UPR^mt^ and mitophagy. These data are consistent with the recently published data that loss of parkin homolog *pdr-1* in *C. elegans* results in up-regulation of *uaDf5* frequency (Valenci, Yonai et al. 2015). Furthermore, overexpression of parkin is reported to result in selection against mutant mtDNA in a mammalian cell line (Suen, Narendra et al. 2010). Taken together, our data suggest that UPR^mt^ ‘protects’ selfish mtDNA like *uaDf5* from organelle level selection.

Many mitochondrial stressors such as exposure to paraquat or disruption of electron transport chain function induce UPR^mt^ (Durieux, Wolff et al. 2011, Runkel, Liu et al. 2013). It will be interesting to determine whether *uaDf5* levels are modulated under these conditions. Severe mitochondrial stressors might sensitize *uaDf5* animals, or trigger additional stress response mechanisms such as mitophagy (Palikaras, Lionaki et al. 2015). In this instance, *uaDf5* levels might decrease due to selection against high levels of *uaDf5* either at the organismal or organelle level. In contrast, *uaDf5* levels might be able to increase under mild mitochondrial stress conditions that activate UPR^mt^ without inducing significant mitophagy or significantly compromising organismal fitness. An elegant series of experiments using RNAi dilution series show that the level of mitochondrial stress is an important determinant of aging (Rea, Ventura et al. 2007). These studies show that mild mitochondrial stress extends lifespan while severe mitochondrial stress can shorten lifespan. Similarly, it will be interesting to test whether varying levels of mitochondrial stress result in differential modulation of *uaDf5* levels, the prediction being that *uaDf5* levels will increase with mild mitochondrial stress but decrease with severe stress.

*uaDf5* mtDNA persists, albeit at lower levels, in *afts-1* loss-of-function mutants even after continuously propagating these animals for five months. These data suggest that first, while UPR^mt^ allows *uaDf5* to rise to high levels, it is not required for *uaDf5* persistence. This interpretation is consistent with our observation that UPR^mt^ activation generally correlates with *uaDf5* levels and is likely not induced at appreciable levels in animals with *uaDf5* below a certain threshold. According to this hypothesis, loss of UPR^mt^ will fail to impact *uaDf5* levels when they drop below this threshold. Second, the persistence of *uaDf5* mtDNA in *atfs-1* loss-of-function mutants suggests roles for additional mechanisms such as exploitation of mtDNA copy number control (see above). In this instance, loss of mtDNA copy number control and UPR^mt^ might be required to force permanent loss of *uaDf5*.

Exploitation of homeostatic processes might underlie persistence of naturally occurring selfish mtDNA. Many natural isolates of *C. briggsae* are heteroplasmic for mutant mtDNA (*nad5*Δ mtDNA) with a ~900bp deletion that disrupts an essential gene (Howe and Denver 2008). This *nad5*Δ mtDNA is widespread in *C. briggsae* natural populations despite its deleterious organismal effects (Estes, Coleman-Hulbert et al. 2011). It has also been shown to have a strong drive to increase in frequency when bottlenecked through small populations (Clark, Howe et al. 2012). It will be interesting to determine in the future whether homeostatic mtDNA copy number control and UPR^mt^ activation contribute to persistence of *nad5*Δ mtDNA in *C. briggsae*.

## Materials & Methods

*Strains*. Worm strains were maintained on nematode growth medium (NGM) seeded with OP50 *E. coli* at 20°C under standard laboratory conditions.

*Mutants*. LGIII, *pdr-1(gk448)*; LGV, *atfs-1(tm4525)*, *atfs-1(et15)*; mtDNA, *uaDf5*, *mptDf1*.

*Transgenic lines. zcIs9 [hsp-60::GFP + lin-15(+)], zcIs17 [ges-1::GFP(mit)], zcIs18 [ges-1::GFP(cyt)]*, *byEx655 [pink-1::Pink-1:GFP + myo-2::mcherry + herring sperm DNA]*.

*Detecting presence of uaDf5 and mptDf1*. To confirm the presence of heteroplasmic *uaDf5* and wildtype mtDNA, worms were individually transferred to PCR tubes containing 50 μL of lysis buffer with 100 μg/mL proteinase K, incubated at −80°C for 10 minutes, incubated at 60°C for 60 minutes, and then at 95°C for 15 minutes to heat-inactivate the proteinase K. The presence of *uaDf5* and wildtype mtDNA copies was confirmed by PCR using the mutant-specific forward primer 5’-CCATCCGTGCTAGAAGACAA-3’ with the wildtype-specific forward primer 5’-TTGGTGTTACAGGGGCAACA-3’ (in the region spanning the deletion), and reverse primer 5’-CTTCTACAGTGCATTGACCTAGTC-3’ common to both mutant and wildtype mtDNA. The *C. elegans* strain harboring the *mptDf1* deletion was maintained in similar manner and the presence of heteroplasmic mtDNA was confirmed using the forward primer 5’-GATTTTTCTGAAGGTGAAAGGGAG-3’ with the reverse primer 5’-CTCCTACTAACCTATTAAGAAATTTTTTCAC-3’, which flank the deletion and generate a product specific to each template.

*Selection for loss of uaDf5*. Every generation, 8-16 individual animals were grown in isolation and PCR was performed on these single individuals after they had progeny to qualitatively determine *uaDf5* levels. 8-16 individuals from the plate with the lowest *uaDf5* levels were picked and the process was repeated until *uaDf5* was no longer detected.

*Sequencing non-uaDf5 mtDNA*. Non-*uaDf5* mtDNA was amplified as three overlapping PCR products. The coordinates of the PCR products and the primers utilized are as follows:

1-5132bp

Forward primer MP236 (CAGTAAATAGTTTAATAAAAATATAGCATTTGGGTTG)

Reverse primer MP255 (CCGTGGCAATATAACCTAGATGTTCTACC)

4997-13311bp

Forward primer MP210 (TTGGTGTTACAGGGGCAACA)

Reverse primer MP256 (GTATGCAAAACATATATTTTTATTAAACTAAAATCCC)

12691-423bp

Forward primer MP230 (GTGGATTAATTTTCTCAAGGGGTGCTG)

Reverse primer MP257 (CCTAAATATCTTCTATAAGTTAATACTGTGGGAG)

The following primers were used to sequence the above PCR products:

1-5132bp PCR product

Reverse primer MP257 (CCTAAATATCTTCTATAAGTTAATACTGTGGGAG)

Forward primer MP237 (GTGTTTTTTGTTTTAGCTGTTTTAAGTAGG)

Forward primer MP249 (GTGTTTTTCTGTTATTTCAAGAATCCTGGG)

Forward primer MP250 (GGAGGCTGAGTAGTAACTGAGAACCC)

Forward primer MP212 (CTTTTATTACTCTATATGAGCGTC)

Forward primer MP209 (CCATCCGTGCTAGAAGACAA)

Reverse primer MP252 (CGCACTGTTAAAGCAAGTGGACGAG)

Forward primer MP251 (GCTGCTGTAGCGTGATTAAGTACTTTG)

Forward primer MP253 (GTTCTAGGTTAAATCCTGCTCGTTTTTG)

Forward primer MP254 (GAGTCTTTTAATTGGATTGTTTTGGGAG)

Reverse primer MP255 (CCGTGGCAATATAACCTAGATGTTCTACC)

4997-13311bp PCR product

Forward primer MP210 (TTGGTGTTACAGGGGCAACA)

Forward primer MP232 (GACTAGGTCAATGCACTGTAGAAGACCC)

Reverse primer MP213 (CGAATTTAAACCCGTCTATAACG)

Forward primer MP240 (TCATCATCTGGGGTTGGAATTTGC)

Forward primer MP241 (CCTAAAGCTCATGTAGAGGCTCCTAC)

Forward primer MP242 (GAAATGTAGGGTTTTCAGCACCATTAGTC)

Forward primer MP243 (CAGCAGGGTTAAGATCTATCTTAGGTGG)

Forward primer MP244 (GGTGGGTTGACAGGTGTTGTATTATC)

Forward primer MP234 (GTCTGTAAGGTTCATACCCTTGAGGTGG)

Reverse primer MP233 (GCCCAAGCATGAATAACATCAGCAGATG)

Forward primer MP245 (CTAGATCAATTAAGTTTAGGTGAACCACG)

Reverse primer MP238 (CTAGCGTAAACACTAAAACTATTAATAGCAC)

Reverse primer MP246 (CTCTAAACGTTACCAAAAAAAGAATAAACG)

Forward primer MP247 (GGTGGAATTAGTGTTTGGCTTATACCCAC)

Forward primer MP248 (GGTGAGGTCTTTGGTTCATAGTAGAAC)

Forward primer MP230 (GTGGATTAATTTTCTCAAGGGGTGCTG)

Reverse primer MP256 (GTATGCAAAACATATATTTTTATTAAACTAAAATCCC)

Reverse primer MP211 (CTTCTACAGTGCATTGACCTAGTC)

12691-423bp PCR product

Reverse primer MP239 (CCACTGCTTAAAAATAAGGTGTACCCC)

Reverse primer MP257 (CCTAAATATCTTCTATAAGTTAATACTGTGGGAG)

Reverse primer MP262 (CAACCCAAATGCTATATTTTTATTAAACTATTTACTG)

Forward primer MP230 (GTGGATTAATTTTCTCAAGGGGTGCTG)

*Droplet digital PCR*. To quantify mtDNA copy number, hermaphrodite worms were selected individually at the fourth and final larval stage (L4) and transferred to OP50-seeded NGM plates. At 96 hours post-transfer, the hermaphrodites were lysed as day 4 adults as described above. Lysates were diluted in nuclease-free water to a factor of 1:50 or 1:100 (the same dilution factor was used across all samples for an experiment) for single-animal lysates and 1:500 or 1:1000 for pooled animal lysates (the same dilution factor was used across all samples for an experiment) and 2 μL of diluted lysate was used in the preparation of each reaction. For ddPCR quantification of wildtype and *uaDf5* mtDNA copy number and heteroplasmy frequency, the forward 5’-GTGATGCAGAGATGTTTATTGAAGC-3’ and reverse 5’-CACTCTGGAACAATATGAACTGGC-3’ primers were used to amplify a wildtype-specific product. The forward 5’-GCGGTATCGTAAGAAAATCAAAATATGG-3’ and reverse 5’-CTTTGTCTTCTAGCACGGATGG-3’ primers were used to amplify a product common to wildtype and mutant templates; *uaDf5* copy number was then determined by subtracting wildtype from total mtDNA copy number. For confirming loss of *uaDf5* (Fig. 1D), the forward 5’-CCATCCGTGCTAGAAGACAAAG-3’ and reverse 5’-CTACAGTGCATTGACCTAGTCATC-3’ primers were used for amplifying a *uaDf5*-specific product. At 297bp, the *uaDf5*-specific product is near the upper size limit for ddPCR; to ensure positive droplet identification, the wildtype-specific and common primer pairs were designed to amplify smaller products (<100bp each) and were used for quantification of *uaDf5* levels. Quantification of *mptDf1* mtDNA (Fig. 2D-2F) was accomplished in a similar manner using L4 animals and the forward 5’-GGATTTAATGTGGAGTTTGCCAGAGTAG-3’ with the reverse 5’-CATAACGATAACGAGGGTATGAACTACG-3’ primers for wildtype-specific amplification, and forward 5’-ATTTCCAATTTATTTTTTACATCTTTGATTACC-3’ with reverse 5’-CCCGCTGTGCCTAATTTTAATAG-3’ for amplification of the common product. For quantification of actin, lysates were diluted 1:5 and the forward primer 5’-CAACACTGTTCTTTCCGGAGG-3’ was used with the reverse primer 5’-GTGATTTCCTTCTGCATACGATC-3’. For specifically detecting *uaDf5*, mutant-specific product was amplified using the forward primer 5’-CCATCCGTGCTAGAAGACAAAG-3’ with the reverse primer 5’-CTACAGTGCATTGACCTAGTCATC-3’. Primers were diluted to 10 μM in nuclease-free water and 0.125 μL of each primer was combined with 2 μL of diluted lysate and 5.5 μL nuclease-free water to bring the reaction volume to 8 μL.

*Quantification of gene expression*. Gene expression was quantified using a commercial cDNA synthesis kit from Thermo Scientific and ddPCR. Lysates were prepared by transferring 10 adult worms to 10 μL lysis buffer with 20 mg/mL proteinase K. The lysates were incubated at 65°C for 10 minutes, 85°C for 1 minute, and 4°C for 2 minutes. RNA transcripts in the worm lysates were immediately converted to cDNA. To accomplish this, 2 μL worm lysate was incubated at 37°C for 2 minutes with 0.5 μL double-stranded DNase (dsDNase), 0.5 μL 10x dsDNase buffer, and 2 μL H_2_O. Following dsDNase incubation, lysates were combined with 0.5 μL oligo d(T) primer, 0.5 μL 10 mM dNTP, 1.5 μL H_2_O, 2 μL 5x reverse transcriptase buffer, 0.5 μL Maxima H Minus reverse transcriptase, and incubated at 25°C for 10 minutes and 55°C for 30 minutes, followed by a 5 minute heat-inactivation at 85°C and 2 minutes at 4°C. The cDNA was then diluted to 50 μL with nuclease-free H_2_O and stored at −80°C. Quantification of cDNA copy number was performed using ddPCR against quadruplicates of cDNA diluted 10-fold in nuclease-free H_2_O. The ddPCR was performed as described above, using primers:

Forward primer MP393 (CAACACTGTTCTTTCCGGAGG) with

Reverse primer MP391 (GTGATTTCCTTCTGCATACGATC) for actin.

Forward primer MP407 (GACGAACACAAACGTGAACGGTTGG) with

Reverse primer MP408 (GGCACTCGGCTGCATACTTTCC) for *nuo-2*.

Forward primer MP-BG-43 (ATTTCCAATTTATTTTTTACATCTTTGATTACC) with

Reverse primer MP-BG-44 (CCCGCTGTGCCTAATTTTAATAG) for *ND4*.

Forward primer MP259 (GAGGTTTTGGTGTTACAGGGGCAAC) with

Reverse primer MP278 (GCATCTTTACCTAAGTACTCAGGTC) for *cytb*.

Forward primer MP481 (GCCATCCGTGCTAGAAGACAAAG) with

Reverse primer MP482 (CCTCTAACTAACTCCCTTTCACCTTCAG) for ND1.

Forward primer MP243 (CAGCAGGGTTAAGATCTATCTTAGGTGG) with

Reverse primer MP411 (CGATCAGTTAACAACATAGTAATAGCCCC) for COXI.

Forward primer MP245 (CTAGATCAATTAAGTTTAGGTGAACCACG) with

Reverse primer MP421 (CCAAGCATGAATAACATCAGCAGATG) for COXII.

Forward primer MP477 (GCTTGAGGTAAGGATATTGCTATAGAAGG) with

Reverse primer MP478 (GTGTACTGGTACTAGAGCAGCATC) for COXIII.

Forward primer MP479 (GATCTTGGTTACCCAAAGCTATAAGAGC) with

Reverse primer MP480 (GTGTCCTCAAGGCTACCACCTTCTTC) for ND5.

Forward primer MP473 (CTTGAGCCATCGTCGATTATTGATG) with

Reverse primer MP474 (CATCTTGGAGAGCTGTGCGAACC) for *hsp-60*.

Forward primer MP475 (GAAGATACGAAGACCCAGAGGTTC) with

Reverse primer MP476 (GAACGAATGCTCCAACCTGAGATG) for *hsp-6*.

*Genetic crosses*. To construct strains with *zcIs9, zcIs17, and zcIs18* heteroplasmic for *uaDf5* mtDNA, animals homozygous for the transgene were selected at the L4 stage, heat-shocked at 30°C for six hours to increase nondisjunction frequency, then stored at 20°C. On a fresh OP50-seeded plate, 7-9 transgenic males from the subsequent generation were combined with 3-5 hermaphrodites heteroplasmic for *uaDf5* mtDNA. Hermaphrodites from the F1 generation were replated at the L4 stage to allow self-fertilization. F2 L4 hermaphrodites were then individually selected and replated. The F2 plates exhibiting presence of GFP fluorescence in 100% of F3 progeny were identified as homozygous for the transgene. Retention of *uaDf5* in the resulting strains homozygous for the transgene was confirmed by PCR. The *uaDf5* heteroplasmy was crossed into a nuclear background harboring the *atfs-1*(*tm4525*) loss-of-function allele using the same protocol as described for the transgenes, with the caveat that F2 plates were identified as homozygous for the *atfs-1*(*tm4525*) loss-of-function allele according loss of GFP fluorescence. The homozygous loss of *atfs-1(tm4525)* was confirmed with

PCR using primers: GAAACCGCCTCCTTTCGCCTTTTG, GACTTCATCGTCGTCCATGGGTACG, and

TCTCCAATTTTGTTAACTTCCAGCAGCC. The *uaDf5* heteroplasmy was crossed into a nuclear background harboring the *atfs-1*(*et15*) gain-of-function allele using the same protocol as described for the *atfs-1(tm4525)* loss-of-function allele; however, ddPCR was conducted on F1 gain-of-function heterozygotes since *atfs-1*(*et15*) homozygous animals are slightly slow growing. The presence of *atfs-1(et15)* allele was confirmed by sequencing. Presence or absence of *pdr-1(gk448)* genotypes were confirmed with PCR using primers: GAATCATGTTGAAAATGTGACGCGAG, CTGACACCTGCAACGTAGGTCAAG, and GATTTGACTAGAACAGAGGTTGACGAG.

*Fluorescence microscopy*. Fluorescence images of worms were captured using Zeiss Axio Zoom V16 stereo zoom microscope. The fluorescence intensities were quantified using Image J. 25 animals were used to calculate average fluorescence intensity for each group. To correlate *hsp60::gfp* fluorescence with *uaDf5* levels, worms heteroplasmic for *uaDf5* mtDNA and expressing the *hsp60::gfp* marker were individually picked as day 4 adults and immobilized on unseeded NGM plates treated with 250 μL 10 mM levamisole. Worms were individually imaged, followed by lysis and ddPCR quantification of mtDNA heteroplasmy as described above.

*TMRE and Mitotracker green FM staining*. 250 μL of 10 μM TMRE dye (Molecular Probes) was added to NGM plates seeded with OP50 and allowed to dry. Adult animals were grown on these plates for overnight. These animals were picked onto new OP50 plates without the TMRE dye and imaged one hour later. Same protocol was followed to stain animals with Mitotracker Green FM (Molecular Probes), but 250ul of 50uM was used.

*RNAi-mediated gene knockdown*. RNAi-mediated gene knockdown was induced using feeder plates. Cultures consisting of 2 mL LB and 10 μL ampicillin were inoculated with bacteria harboring the ZC376.7 (*atfs-1*) ORF plasmid clone and incubated on a shaker at 37°C overnight. Next, 750 μL of each overnight culture was transferred to a flask containing 75 mL LB and 375 μL ampicillin. The 75 mL culture was incubated on a shaker at 37°C for 4-6 hours, until OD_550-600_ > 0.8. An additional 75 mL LB was added to the culture along with 375 μL ampicillin and 600 μL 1 M isopropyl β-D-1-thiogalactopyranoside (IPTG) to induce expression of the small interfering RNA (siRNA). Cultures were incubated an additional 3.5-4 hours on a shaker at 37°C and centrifuged at 3900 rpm for 6 minutes. After discarding the supernatant, the resulting pellets were resuspended in 6 mL M9 buffer with 8 mM IPTG and 250 μL of resuspension was pipetted onto unseeded NGM plates. Once dry, RNAi feeder plates were stored at 4°C. Control plates were prepared using bacteria harboring an empty (no siRNA) vector. Worms were transferred at the L4 stage to empty vector control plates and RNAi plates and stored at 20°C. Additional L4 worms were selected from the F1 generation, aged to day 4 adults on fresh RNAi plates, and lysed as described above. For multigenerational RNAi, L4 worms were transferred to fresh RNAi and empty vector control plates every 2-3 days, and lysed after seven generations.

*Fitness assays*. Three adults picked from population of animals growing on RNAi plates since L4 stage were allowed to lay eggs for 3 hrs on corresponding fresh RNAi plates. Number of unhatched and hatched embryos were counted one day later to determine percentage of unhatched embryos. After additional two days, total number of larva and adults were counted to determine fraction of animals that experienced delayed growth. Subsequently, all animals were transferred to fresh RNAi plates every other day until day four of adulthood. Number of total dead animals were counted until day four of adulthood to determine percentage of dead animals.

*Western blot analysis*. One hundred adult worms in 10 μL M9 were lysed in 10 μL of 2X SDS sample buffer and boiled for 10 minutes. SDS PAGE gel and transfer were performed according to standard protocol. Mouse monoclonal anti-beta-actin (sc-47778, Santa Cruz Biotechnology) or mouse monoclonal anti-GFP (sc-9996, Santa Cruz Biotechnology) were used at 1:500 dilution overnight at 4°C as primary antibodies. HRP-conjugated goat anti-mouse antibody (sc-2005, Santa Cruz Biotechnology) was used at 1:5000 dilution for 90 minutes at room temperature as the secondary antibody. SuperSignal West Pico Chemiluminescent Substrate (Thermo Fisher) was used for detecting HRP.

## Author contributions

MRP and BLG conceived and designed the study. BLG and MRP carried out the experiments. CSK carried out experiments with *mptDf1* mtDNA heteroplasmy. BLG and RDG performed ddPCR experiments in the laboratory of SAM. MRP, BLG, and DCS analyzed the data. BLG and MRP prepared the manuscript.

## Acknowledgements

*uaDf5* and *mptDf1* heteroplasmic strains, pdr-1(gk448) and *atfs-1(e15)* mutant strain, and *zcIs9*, *zcIs17*, *zcIs18*, and *byEx655* transgenic lines were kindly provided by Caenorhabditis Genetics Center (CGC). *atfs-1(tm4525)* mutant strain was kindly provided by the Mitani Lab through the National Bio-Resource Project of the MEXT, Japan. We would like to thank Sarah Sturgeon for technical assistance. We would like to thank O. Thompson and R. Waterston (University of Washington) for identifying *mptDf1* deletion from the Million Mutation Project worm collection. We would like to thank Harmit Malik (HHMI/Fred Hutchinson Cancer Research Center) for valuable advice and support. We thank Harmit Malik, Nitin Phadnis, and Janet Young for providing critical comments on the manuscript. Funding for this work was provided in part by Helen Hay Whitney Foundation Fellowship (MRP), by grants from the Mathers Foundation and HHMI (to Harmit Malik), startup funds from Vanderbilt University (MRP), and the NIH-funded Tennessee Center for AIDS Research (P30 AI110527) (to SAM).

